# Nonlinear transcriptional responses to gradual modulation of transcription factor dosage

**DOI:** 10.1101/2024.03.01.582837

**Authors:** Júlia Domingo, Mariia Minaeva, John A Morris, Samuel Ghatan, Marcello Ziosi, Neville E Sanjana, Tuuli Lappalainen

## Abstract

Genomic loci associated with common traits and diseases are typically non-coding and likely impact gene expression, sometimes coinciding with rare loss-of-function variants in the target gene. However, our understanding of how gradual changes in gene dosage affect molecular, cellular, and organismal traits is currently limited. To address this gap, we induced gradual changes in gene expression of four genes using CRISPR activation and inactivation. Downstream transcriptional consequences of dosage modulation of three master trans-regulators associated with blood cell traits (*GFI1B*, *NFE2*, and *MYB*) were examined using targeted single-cell multimodal sequencing. We showed that guide tiling around the TSS is the most effective way to modulate *cis* gene expression across a wide range of fold-changes, with further effects from chromatin accessibility and histone marks that differ between the inhibition and activation systems. Our single-cell data allowed us to precisely detect subtle to large gene expression changes in dozens of *trans* genes, revealing that many responses to dosage changes of these three TFs are nonlinear, including non-monotonic behaviours, even when constraining the fold-changes of the master regulators to a copy number gain or loss. We found that the dosage properties are linked to gene constraint and that some of these nonlinear responses are enriched for disease and GWAS genes. Overall, our study provides a straightforward and scalable method to precisely modulate gene expression and gain insights into its downstream consequences at high resolution.

## Introduction

Precision control of gene expression levels plays a pivotal role in defining cell type specificity and coordinating responses to external stimuli. Imbalances in this intricate regulation can underlie the genetic basis of both common and rare human diseases. The vast majority of genetic variants associated with complex disease, as revealed by genome-wide association studies (GWAS), are located in noncoding regions, with likely gene regulatory effects ^1^. Previous studies have attempted to elucidate these effects by mapping genetic associations to gene expression ^2,3^, and more recently, CRISPR-based perturbations of GWAS loci have provided insights into their functional consequences ^4^. A major driver of rare genetic diseases is loss-of-function variants affecting one or both copies of the gene, leading to disease via dramatic reduction of functional gene dosage ^5^. The substantial overlap ^6,7^ and potential joint effects ^8,9^ of rare and common variants indicate a general link between different degrees of perturbation of gene dosage and disease phenotypes.

However, our understanding of the quantitative relationship between gradual changes in gene dosage and downstream phenotypes remains elusive for most human genes. Practical applications of the compelling allelic series concept to identify genes where increasingly deleterious mutations have increasing phenotypic effects have been limited by the sparsity of segregating variants with an impact on a given gene in the human population ^10^. Experimental characterization of gene function in model systems has predominantly relied on gene knock-out or knock-down approaches ^11^. While these studies have proven useful to identify dosage-sensitive genes involved in cellular functions and disease ^12–16^, these approaches only provide a limited discrete relationship between the number of functional gene copies and a certain phenotype (eg. loss-of-function consequence vs. wild-type). However, such relationships are in fact determined by continuous dosage-to-phenotypes functions that, as suggested by a small number of previous experimental studies ^17–19^, can be complex and thus are challenging to infer from loss-/gain-of-function data.

Recently, new methods have enabled the gradual modulation of gene dosage in model systems ^18,20–22^, while large-scale insights into the downstream effects of dosage modulation have largely come from yeast ^17^ and bacteria ^19,23^, demonstrating that nonlinear relationships between gene dosage and phenotype are common. In humans, the relationship between dosage and downstream phenotypes is largely unexplored. Only a few limited studies ^17–19^ have dissected these consequences. For instance, the disease-associated transcription factor *SOX9* ^24^ showed a nonlinear relationship between dosage and multiple tiers of phenotypes, including DNA accessibility, RNA expression of downstream targets, raising the question of whether this phenomenon occurs with other transcription factors. More recently, similar evidence has been shown in the case of the *NKX2-1* lineage factor with an oncogenic role in lung adenocarcinoma ^25^. Generally, transcription factors represent a particularly compelling target for the characterization of gene dosage effects. They are key regulators of cellular functions, enriched for disease associations ^26^ and often classified as haploinsufficient ^27^. Additionally, their effects can be measured by transcriptome analysis. However, our knowledge of their dosage-dependent effects on regulatory networks still remains limited, particularly regarding subtle dosage variation within their natural range ^22^.

In this study, we developed and characterised a scalable novel sgRNA design approach for gradually decreasing and increasing gene dosage with the CRISPR interference (CRISPRi) and activation (CRISPRa) systems. We applied this to four genes, with single-cell RNA-sequencing (scRNA-seq) as a cellular readout of downstream effects. While classic Perturb-Seq analyses have focused on gene knockdown effects, we assess the effects of gradual up- and down-regulation of target genes. We uncovered quantitative patterns of how gradual changes in transcription dosage lead to linear and nonlinear responses in downstream genes. Many downstream genes are associated with rare and complex diseases, with potential effects on cellular phenotypes.

## Results

### Precise modulation and quantification of gene dosage using CRISPR and targeted multimodal single-cell sequencing

We selected four genes for gradual modulation of their dosage in the human erythroid progenitor cell line K562 ^28^: *GFI1B*, *NFE2*, *MYB* and *TET2*. Two of the genes, *GFI1B* and *NFE2*, have been implicated in blood diseases and traits ^29–31^, and in our earlier work, we identified a broad transcriptional response to inhibition of GWAS-overlapping enhancers to these genes ^4^. *MYB* is a key transcription factor ^32^ and a downstream target of *GFI1B* ^4^.*TET2* has a role in DNA demethylation and is unrelated to these transcriptional networks and is included in this study as a control with minimal expected *trans* effects. We refer to these four genes, targeted in *cis* for modulation of their regulation, as *cis* genes (**Figure 1A**).

**Figure 1:**
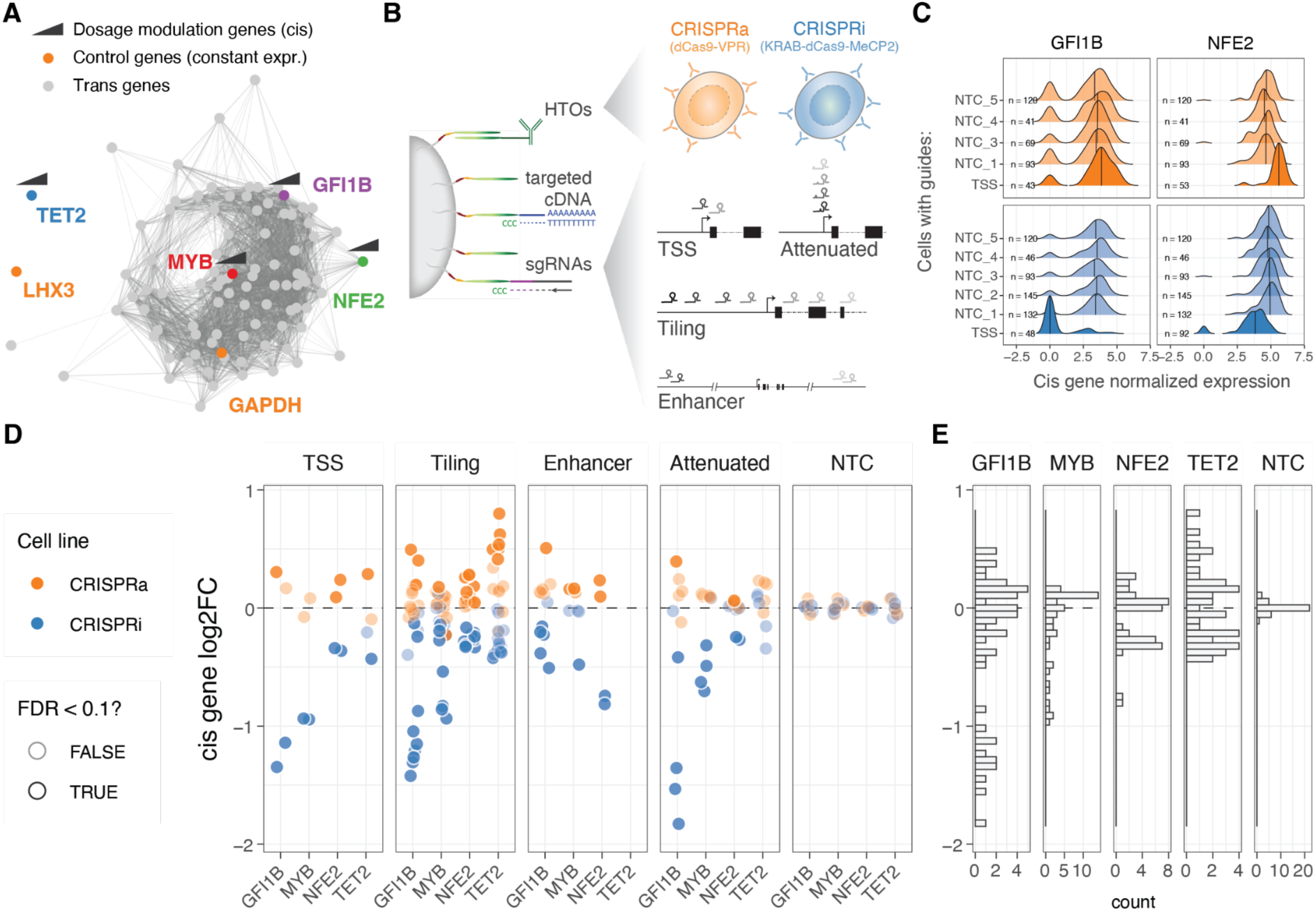
Modulation and quantification of gene dosage using CRISPR and targeted multimodal single-cell sequencing. A. Co-expression network representation of the 92 selected genes under study. Genes (nodes) are connected by edges when their co-expression across single cells was above 0.5 (data used from Morris *et al*. 2023). Highlighted in colour are the two control highly (GAPDH) and lowly (LHX3) constantly expressed genes, as well as cis genes for which dosage was modulated with CRISPRi/a. B. Design of the multimodal single cell experiment (HTO = hash-tag oligos). C. Distribution of the GFI1B (left) or NFE2 (right) normalised expression across single cells for different classes of sgRNAs (NTC = Non-targeting controls, TSS = transcription start site). D. Resulting relative expression change (log2 fold change) of the 4 cis genes upon each unique CRISPR perturbation when grouped across different classes of sgRNAs. E. Distribution of cis gene log2FC across all sgRNA perturbations.

To modulate the gene expression of the *cis* genes, we use K562 cells expressing CRISPRi (KRAB-dCas9-MECP2) and CRISPRa (dCas9-VPR) systems (see Methods), both cell lines were hashed with DNA conjugated antibodies against different surface proteins that allow pooled experiments. To obtain a wide range of dosage effects we used four different single guide RNA (sgRNA) design strategies (**Figure 1B**): 1) targeting the transcription start site (TSS) as in the standard CRISPRi/CRISPRa approach, 2) tiling sgRNAs +/- 1000 bp from the TSS in approximately 100bp intervals, 3) targeting known *cis*-regulatory elements (CREs), and 4) using attenuated guides that target the TSS but contain mismatches to modulate their activity ^18^. We further included 5 non-targeting control (NTC) sgRNAs as negative controls.

The library of altogether 96 guides was transduced to a pool of K562-CRISPRi and K562-CRISPRa cells at low multiplicity of infection (MOI). After eight days, we performed ECCITE-seq (see Methods) to capture three modalities: cDNA, sgRNAs and surface protein hashes (oligo-tagged antibodies with unique barcodes against ubiquitously expressed surface proteins). Instead of sequencing the full transcriptome, we used target hybridization to capture a smaller fraction of the cDNA and obtain more accurate expression readouts at a feasible cost. The subset of selected transcripts were picked from the transcriptional downstream regulatory networks of *GFI1B* and *NFE2* identified previously ^4^, maintaining similar patterns of co-expression correlation across co-expression clusters (see Methods, **Figure S1A**). We targeted a total of 94 transcripts (**Figure 1A**), including the four *cis* genes, 86 genes that represent trans targets of *GFI1B* and/or *NFE2* ^4^ (**Figure S1A**), *LXH3* that is not expressed in blood progenitors, *GAPDH* that is highly expressed and often considered an invariable housekeeping gene and the dCas9-VPR or KRAB-dCas9-MeCP2 transcripts.

We used the protein hashes and the dCas9 cDNA (indicating the presence or absence of the KRAB domain) to demultiplex and determine the cell line—CRISPRi or CRISPRa. Cells containing a single sgRNA were identified using a Gaussian mixture model (see Methods). Standard quality control procedures were applied to the scRNA-seq data (see Methods). To confirm that the targeted transcript capture approach worked as intended, we assessed concordance across capture lanes (Figure S1C). The final data set had 20,001 cells (10,647 CRISPRi and 9,354 CRISPRa), with an average of 81 and 86 cells with a unique sgRNA for the CRISPRa and CRISPRi, respectively (**Figure S1D**).

### Gradual modulation of gene expression across a broad range with CRISPRi/a

Next, we calculated the expression fold change for each of the four *cis* genes targeted by each sgRNA in the two cell lines (CRISPRi/a), comparing each group of cells with its respective NTC sgRNA group (see Methods). We first confirmed that the sgRNAs targeting the transcription start site (TSS) up- and down-regulated their targets (**Figure 1C**, **Figure S1F**). When looking at all sgRNAs at once, across the four genes, we observed a 2.3 fold range (**Figure 1E**), with a minimum 72% reduction and maximum 174% increased expression (log2(FC) values from -1.83 to 0.80). However, the range varied between the genes, with *GFI1B* covering the widest range of gene expression changes (gene expression ranging between 0.28 to 1.42 fold), while *MYB* expression could not be pushed higher than 1.13 fold (**Figure 1E**). The direction of the effects were consistent with the cell lines of origin, where 98.88% of the significant perturbations (Wilcoxon rank test at 10% FDR, n = 89) were correctly predicted based on the direction of the target gene fold change. The predicted on- and off-target properties of the guides ^33–35^ did not correlate with the fold changes in the *cis* genes (**Figure S2A**), suggesting that the observed effects represent true *cis*-regulatory changes. The fold changes were also robust to the number of cells containing a particular sgRNA (**Figure S2B**, top).

We verified that the fold change estimation was not biased depending on the expression level of the target gene at the single-cell level, which can vary due to drop-out effects or binary on/off effects of the KRAB-based CRISPRi system ^20^. By splitting cells with the same sgRNA based on the normalised expression of the *cis* gene (0 vs. >0 normalised UMIs, **Figure S3A**), we observed highly concordant transcriptome gene expression effects between the two groups (**Figure S3B**). This indicates that the dosage changes per guide were not primarily driven by the changing frequency of binary on/off effects, and the use of pseudo-bulk fold changes provides a robust estimation of *cis* gene fold changes. These patterns are further supported by the cells forming a gradient rather than distinct clusters on a UMAP (**Figure S3C**).

The fold change patterns differed between sgRNA designs (**Figure 1D**, left). As expected, sgRNAs targeting the TSS showed strong perturbations in gene expression. However, sgRNAs tiled +/- 1kb from the TSS provided a broader and more gradual range of up- and downregulation across the target genes, sometimes surpassing the effects of TSS-targeting sgRNAs. Attenuated sgRNAs with mismatch mutations resulted in a range of gene silencing effects in the CRISPRi line, as expected based on their original design ^18^. However, these attenuated sgRNAs did not exhibit such a dynamic range in the CRISPRa modality, although a significant correlation existed between the silencing or activating effect size and the distance of the mismatch from the protospacer adjacent motif (PAM) when considering all data points together (**Figure S2C**). The sgRNAs targeting distal *cis*-regulatory elements (CREs) showed both inhibiting and activating effects, even though both the CRISPRi and CRISPRa constructs were initially designed to inhibit or activate transcription from the promoter and initial gene body region. Nonetheless, the number of known CREs per gene is typically limited. Given its simplicity and the ability to achieve both up- and downregulation of the target gene, we consider the tiling sgRNA approach, with a simple design that only requires annotation of the TSS, as a useful method for gradually modulating gene dosage with CRISPRi/a systems.

### *Cis* determinants of dosage

Having designed guides targeting both distal and local neighbouring regulatory regions of the four transcription factors (TFs) and ensuring minimal bias in fold-changes due to sgRNA’s biochemical properties, we investigated the *cis* features that determine the strength of dosage perturbation. We observed substantial differences in the effects of the same guide on the CRISPRi and CRISPRa backgrounds, with no significant correlation between *cis* gene fold-changes (**Figure 2A**). However, in both modalities, the strongest effects on gene expression were observed when the guides were close to the transcription start site (TSS) (**Figure 2B**, excluding NTC and attenuated sgRNAs), although the peaks of strongest activation or repression differed between the modalities. In the CRISPRi modality, the maximum effect was located within the gene body at +238 bp from the TSS (**Figure 2B**, bottom), consistent with previous studies that used essentiality as a proxy for expression ^36^. However, in the CRISPRa modality, the maximum average fold changes occurred closer to the TSS at around -99 bp (**Figure 2B**, bottom), as also shown for CD45 ^37^.

**Figure 2:**
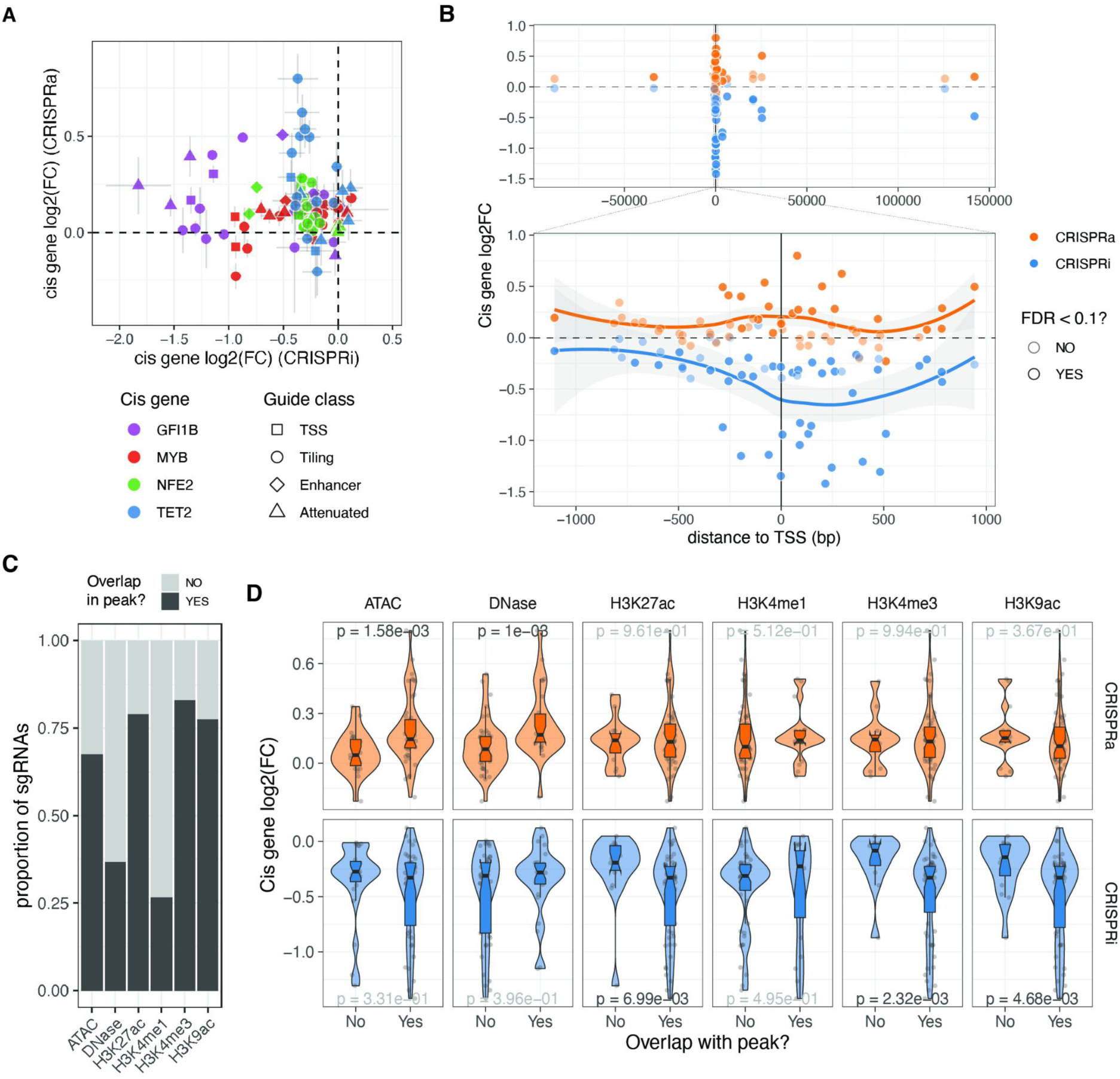
Cis determinants of dosage. A. Comparison of the relative expression change (log2FC) from the same sgRNA between the two different CRISPR modalities. Vertical and horizontal bars represent CRISPRa and CRISPRi standard errors, respectively. B. Relative expression change of the targeted cis gene based on distance from transcription start site (TSS). Top plot excluded attenuated and NTC sgRNAs, while bottom plot also excludes enhancer sgRNAs. C. Number of sgRNAs that overlap with the different epigenetic or open chromatin peaks. D. Relative expression change to NTC sgRNAs (log2(FC)) of all cis genes when their sgRNAs fall or not in the different epigenetic or open chromatin peaks. P-value result from Wilcoxon rank-sum tests, with nominally significant p-values shown in black.

Enhancer, tiling, and TSS sgRNAs targeted regions of the genome with different compositions of histone marks, annotated by ENCODE, in K562 cells^38^ (**Figure 2C**), which allowed us to investigate the impact of chromatin state on the strength of *cis* gene dosage modulation. The magnitude of *cis* gene fold changes varied significantly depending on the presence of specific marks or peaks, which again differed between the two modalities (**Figure 2D**). In the CRISPRa cell line, the strongest effects were observed when guides were located in regions with open chromatin marks, such as DNase or ATAC peaks. In contrast, the strongest repression by CRISPRi occurred in genomic regions with the presence of H3K27ac, H3K4me3, and H3K9ac marks. These differences may be explained by the distinct mechanisms of action of the activator and repressor domains. MeCP2 and KRAB repressor domains recruit additional repressors that silence gene expression through chromatin remodelling activities such as histone deacetylation ^39^. On the other hand, the VPR activation fusion domain may only require Cas9 to scan the open chromatin and recruit RNA polymerase and additional transcription factors to activate transcription. Overall, while a few sgRNAs have a strong effect in both CRISPRi and CRISPRa cell lines, a single guide library containing guides optimised for both modalities enables a range of gradual dosage regulation. However, larger data sets are needed for more careful modelling of the ideal dosage modulation designs and to understand how both *cis*-regulatory features, feedback loops, and other mechanisms contribute to the outcomes.

### *Trans* responses of transcription factor dosage modulation

We then turned our attention to the remaining 91 genes captured by our custom panel and determined the relative expression fold change of each *trans* gene, compared to NTC in each unique guide perturbation (see Methods). Principal component analysis (PCA) performed on all pseudo-bulk fold changes demonstrated the removal of batch effects from the cell lines and revealed a clear direction of the *cis* gene dosage effect in the first three principal components (**Figure S4B**). This finding suggests that dosage modulation is the primary determinant of *trans* effects. The PCA indicated that the dosage modulation of *GFI1B* and *MYB* is reflected in opposite directions in PC1 and PC2, while the *trans* responses of *NFE2* are captured by PC3.

Using a false discovery rate (FDR) cutoff of 0.05, all 91 *trans* genes except for the neural-specific TF *LHX3* (negative control) exhibited a significant change in expression upon perturbation of any of the TFs. The observed trans-effects were well correlated with perturbations of these genes in other data sets (**Figure S4C,D**). Among all measured fold changes, the most extreme negative effect sizes were observed in *cis* genes, with the top 10 being predominantly reductions in *GFI1B* expression. This indicates that *cis* downregulation tended to surpass the endogenous expression limits. In contrast, the largest increases in gene expression were observed through *trans* mechanisms, where *KLK1* and *TUBB1* reached the largest expression values when *GFI1B* was strongly upregulated, or *SPI1* and *DAPK1* when *GFI1B* was strongly downregulated. These findings suggest that the CRISPRa approach did not reach a biological ceiling of overexpression.

Inspecting *trans* responses as a function of *cis* gene modulation, we observed that the number of expressed genes and the mean absolute expression changes of *trans* genes exhibited gene-specific correlations with *cis*-gene dosage (**Figure 3A**, **Figure S4E**). Perturbations in *GFI1B* led to the most pronounced *trans* responses, with positive dosage changes resulting in larger effect sizes compared to decreasing TF gene expression, where the effect plateaued. *NFE2* exhibited similar patterns but with a smaller magnitude. In the case of *MYB*, *trans* responses were observed when decreasing the expression of this TF, but the effects of upregulation are largely unknown as we were unable to increase *MYB* expression beyond 0.35. As expected, given the unrelatedness of *TET2* to the *trans* network, dosage modulation of this gene had minimal *trans* effects with the least pronounced trend when compared to *TET2* dosage, so we excluded it from subsequent analyses.

**Figure 3:**
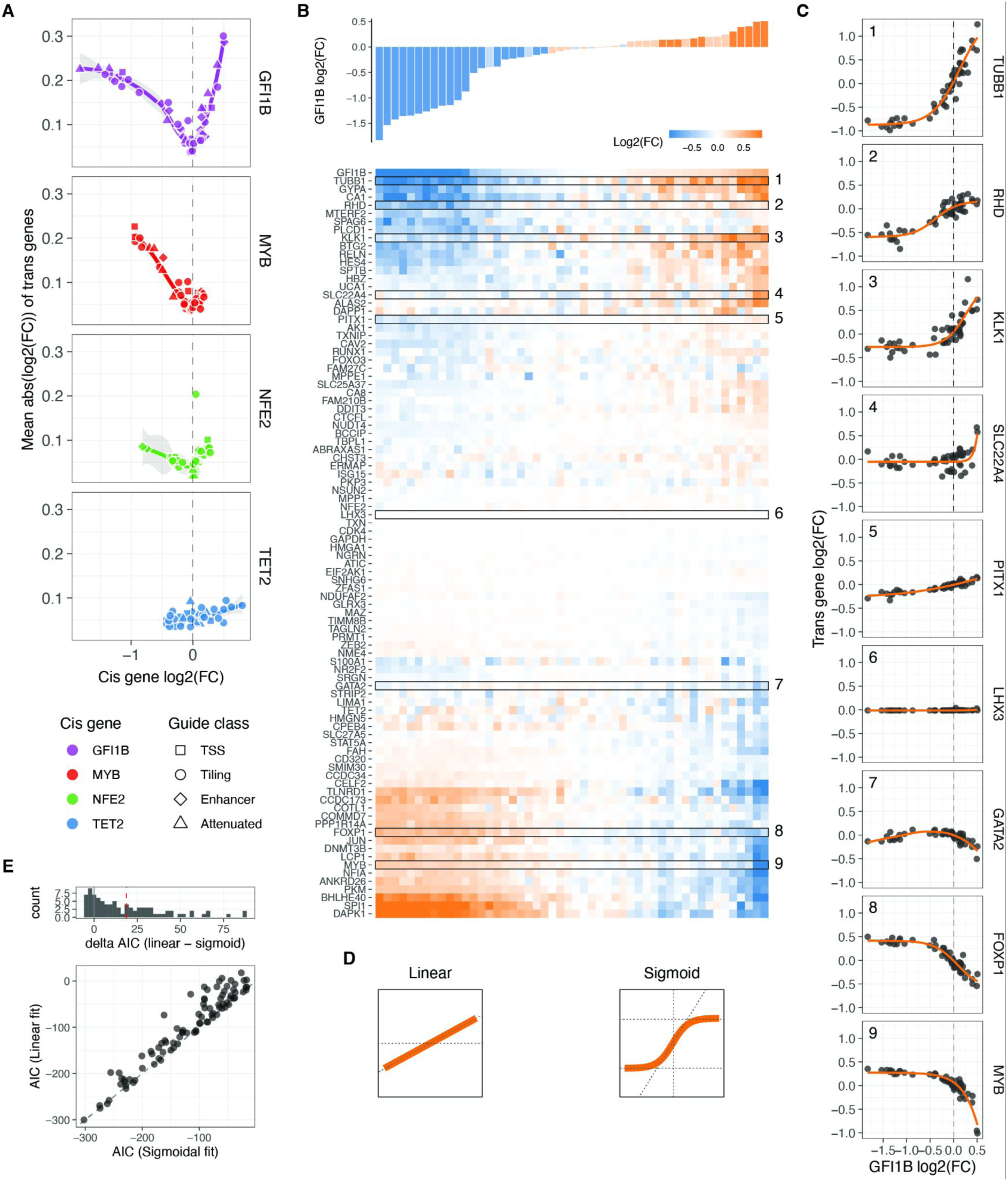
Trans responses of transcription factor dosage modulation. A. Average absolute expression change of all trans genes relative to the changes in expression of the cis genes. B. Changes in relative expression of all trans genes (bottom heatmap) in response to GFI1B expression changes (top barplot) upon each distinct targeted sgRNA perturbation, in comparison to NTC cells. The rows of the heatmap (trans genes) are hierarchically clustered based on their expression fold change linked to alterations in GFI1B dosage. Highlighted rows are selected dosage response examples shown in panel C. C. Dosage response curves of the highlighted trans gene in B as a function of changes in GFI1B expression. The orange line represents the sigmoid model fit, except for GATA2, which display a non-monotonic response and are fitted with a loess curve. D. Illustration of the linear and sigmoid models and equations used to fit the dosage response curves. E. Distribution of the difference in Akaike Information Criterion (ΔAIC_linear-sigmoid_) after fitting the sigmoidal or linear model for each trans gene upon GFI1B dosage modulation (top panel), and the direct comparison of the AIC of each fit (bottom panel).

### Widespread nonlinear dosage responses in *trans* regulatory networks

Upon clustering the changes in expression of *trans* genes based on the *cis* gene dosage change linked to each sgRNA, we identified distinct clusters exhibiting different dosage-response patterns (**Figure 3B** for *GFI1B*, **Figure S5-8A** for all *cis* genes). Further examination of the gene expression fold changes for each individual trans gene in relation to the TF fold changes revealed a diverse range of response patterns (**Figure 3C**, **Figure S5-8B** for all *cis* genes). These responses exhibited both linear and nonlinear forms, including some instances of non-monotonic gene expression responses for certain *trans* genes within the *GFI1B* trans network (e.g., *GATA2* in **Figure 3C**, **Figure S9E**).

To accurately characterise the dosage response, we employed both linear and nonlinear modelling approaches (**Figure 3D**), which allowed us to quantitatively assess the extent of nonlinear responses by comparing the goodness of fit of these models using the Akaike Information Criterion (AIC). For the nonlinear model, we utilised a sigmoid function with four free parameters (**Figure 3D**, right). These parameters represented the slope at the inflection point (*b*, indicating the rate of increase or decrease in expression), the minimum and maximum asymptotes (*c* and *d*, representing the lower and upper limits of fold change), and the value of *cis* gene expression at which the inflection point occurs (*a*). To prevent overfitting, we implemented a 10-fold cross-validation scheme, which yielded reliable predictions on the left-out data (Pearson r = 0.71 to 0.88 for all *trans* genes in the *GFI1B*, *MYB*, and *NFE2* networks, **Figure S9C**). Additionally, the predicted parameter *a* was centred around zero, as expected since the input data represents relative fold changes (**Figure S10**). Since a sigmoid function cannot capture non-monotonic responses, we employed a loess regression as an alternative approach for the few genes that exhibited non-monotonic responses (see Methods, **Figure S9D, E**). For the vast majority of genes, the sigmoid (or loess) fit was remarkably good, partially due to the low level of noise in the targeted scRNA-seq data.

We compared the performance of the linear vs. nonlinear models with the ΔAIC (AlC_linear_ - AlC_r_), where a positive ΔAIC means that the sigmoid model captures the variance better in the dosage response than in the linear model. This showed that most *GFI1B*-dependent dosage expression responses are better fit by the sigmoid model (median ΔAIC = 18.7, with 70.4% of all *trans* genes with a significant response having ΔAIC >2, **Figure 3D**). The responses to dosage modulation of *MYB* and *NFE2* were also better captured by the nonlinearities, but to a lesser extent (0.14 and 3.4 median ΔAIC, with 20.8% and 40.7% of all

*trans* genes dosage responses having ΔAIC > 2 for *MYB* and *NFE2*, respectively, **Figure S9A**). The broader range of *GFI1B* expression modulation, providing more data to detect nonlinear trends, likely contributes to this difference. When ignoring those genes classified as unresponsive (genes that their expression did not change upon the TF modulator, see Methods), even more responses of the remaining *trans* genes were better explained by a sigmoidal model with 83.6%, 26.3% and 63.2% of these having a ΔAIC > 2, for *GFI1B*, *MYB* and *NFE2* respectively. A similar trend holds even when limiting the models to be fitted to those data points that correspond to a hypothetical one copy loss or gain of the *cis* gene (**Figure S9B**), where the median ΔAIC of responsive genes are 7.05, 0.05, and 3.6 for *GFI1B*, *MYB,* and *NFE2 trans* responses. Overall, this shows that *trans* responses to TF dosage are dominated by nonlinear behaviours even when the TF dosage changes are not extreme but within biologically plausible ranges.

### Gene and transcriptional network properties of dosage response

Utilising a model that effectively captures the variance in our data provided the ability to predict unmeasured TF dosage points and facilitated a direct comparison of *trans* effects across different cis genes. Employing the sigmoid model (and loess for those with non-monotonic responses), we estimated the continuous expression of *trans* genes on a uniform fold-change scale across the spectrum of *GFI1B*, *MYB*, and *NFE2* expression changes (**Figure 4A**). This estimation was carried out within the empirically observed range of all three *cis* genes, spanning from log2(FC) -1.83 to 0.51. Subsequent hierarchical clustering of *trans* gene responses revealed six major clusters of distinct response patterns. For the majority of *trans* genes, the response to *GFI1B* and *MYB* was opposite, with only two small clusters displaying exceptions. Notably, *GFI1B* generally induced the most substantial response, while *NFE2* triggered the smallest range of *trans* gene response.

**Figure 4:**
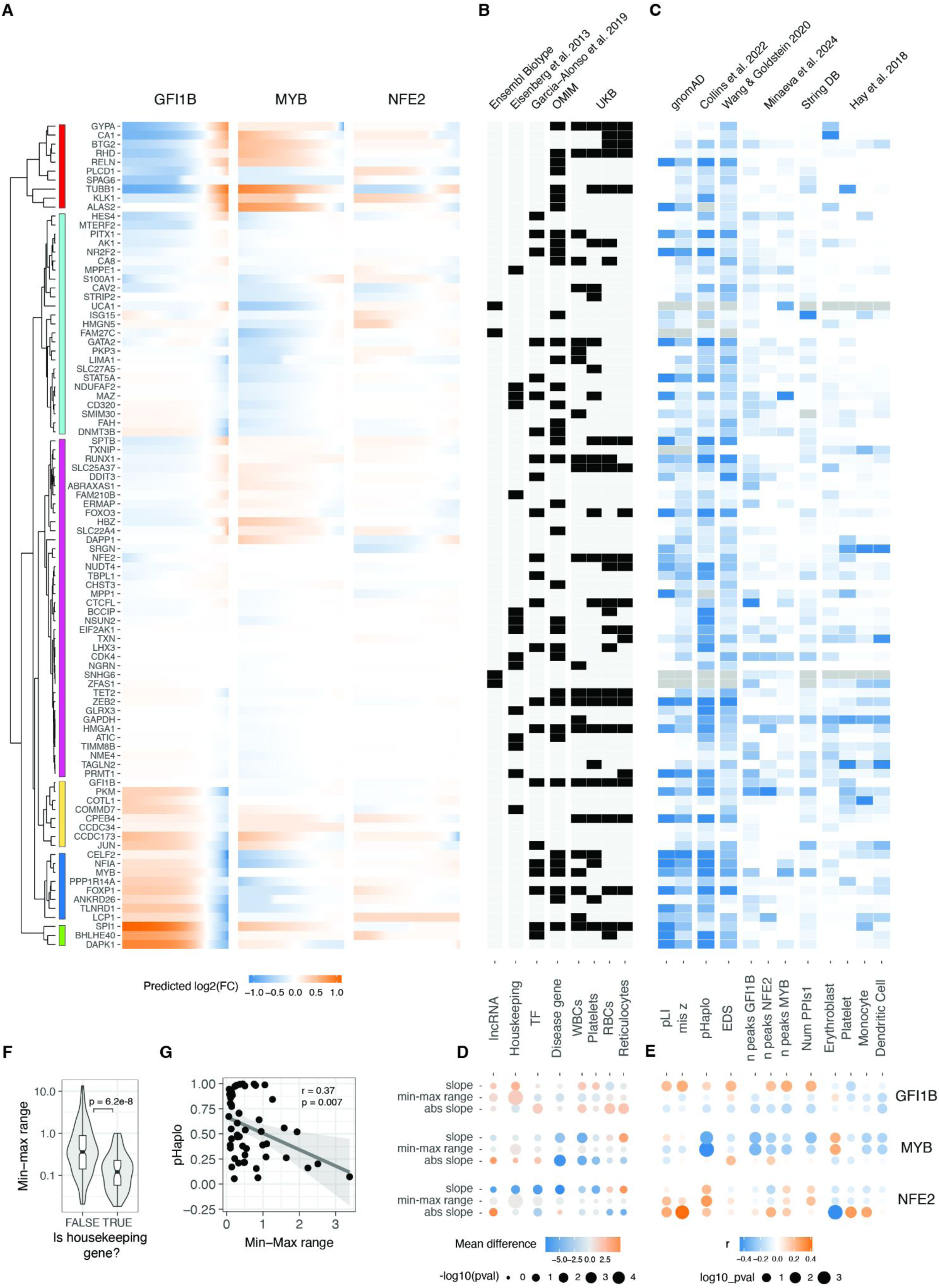
Relationship between gene and dosage response properties. A. Predicted changes (using sigmoid or loess fits for monotonic and non-monotonic responses, respectively) in relative expression of all trans genes in response to changes of the GFI1B, MYB and NFE2 expression. Trans genes (rows) were hierarchically clustered based on their expression fold change linked to alterations of all TF’s dosage. Dendrogram of the resulting clustering shown in the left. B. Heatmap showing the qualitative properties of each trans gene. The x-axis indicates specific gene features. The top labels specify the source of the data, while the bottom labels describe the corresponding gene properties. WBCs, platelets, RBCs, and reticulocytes refer to GWAS of white blood cells, platelets, red blood cells, and reticulocytes, respectively. C. Heatmap indicating the z-scaled quantitative gene features of each transgene. The x-axis indicates specific gene features. The top labels specify the source of the data, while the bottom labels describe the corresponding gene properties. Erythroblast, platelets, monocytes, and dendritic cells refer to cell types from Hay et al. 2018. Grey cells indicate missing data. D. Difference in the average value of the sigmoid parameter indicated in right between the genes qualified into the no/yes category of the gene properties indicated in B. E. Pearson correlation coefficient of the quantitative trans gene features (shown in C) with the sigmoid parameter value for each trans gene in the response of the modulation of dosage of the TF indicated on the left. Size of the points are inversely related to significance of correlation, and colour indicates the direction of correlation. F. Differences in the range of expression response for Housekeeping vs. non-Housekeeping trans genes with changes of dosage of MYB, GFI1B and NFE2. G. Negative correlation between haploinsufficiency score (pHaplo) and the range of the response of trans genes to the modulation of MYB.

Next, we collected diverse annotations for the *trans* genes to explore the connections between their regulatory properties, disease associations, and selective constraints concerning their response to TF dosage (**Figure 4B, C**). To quantify these relationships, we assessed significant differences in belonging to these qualitative annotations using the Wilcoxon rank test (**Figure 4D**) and correlated parameters from the sigmoid model with quantitative gene metrics (**Figure 4E**). We hypothesised that genes with annotated selective constraint, numerous regulatory elements, and central positions in regulatory networks would exhibit greater robustness to TF changes. Indeed, housekeeping genes demonstrated a considerably smaller dosage response range (**Figure 4F**). Moreover, genes classified as unresponsive were enriched in the housekeeping category (odds ratio = 2.14, Fisher test p-value = 0.024). The link between selective constraint and response properties is most apparent in the *MYB* trans network. Specifically, the probability of haploinsufficiency (pHaplo) shows a significant negative correlation with the dynamic range of transcriptional responses (**Figure 4G**): genes under stronger constraint (higher pHaplo) display smaller dynamic ranges, indicating that dosage-sensitive genes are more tightly buffered against changes in *MYB* levels. This pattern was not reproduced in the other trans networks (**Figure 4E**).

The relationship between the response of *trans* genes and intrinsic gene properties differed between *GFI1B*, *MYB,* and *NFE2 trans* network responses. We also performed a similar analysis comparing the sigmoid parameters to network properties using the approach outlined by Minaeva et al.^40^and obtained inconsistent results between TF regulons (**Figure S11A, B**). This suggests that the link between commonly annotated gene properties and the gene responses are complex and highly context specific, as in our data from a single cell line, they differed between the upstream regulators that were manipulated. Thus, much more data is needed before transcriptional responses can be predicted from gene properties, and conversely, to understand the cellular mechanisms that lead to the annotated gene properties.

### Nonlinear dosage responses in complex traits and disease

Moving beyond the characterization of mechanisms of transcriptome regulation, a key question is how gradual dosage variation links to downstream cellular phenotypes, and whether these responses exhibit analogous nonlinear patterns. To address this question, we correlated our findings with the expression profiles of various cell types in order to study the myeloid differentiation process, a phenotype well-characterised for our K562 model that has been used as a reliable system for investigating erythroid differentiation within myeloid lineages ^41^ and blood tumours ^42^. Specifically, leveraging single-cell expression data for bone marrow cell types from the Human Cell Atlas and Human Biomolecular Atlas Project ^43^, we filtered the expression data to the targeted genes in our study. After aggregating data across donors and normalising expression across cell types (**Figure S12A**), we compared the expression patterns resulting from each unique transcription factor dosage modulation in relation to each unique cell type expression state. The ensuing correlation can then be construed as a "phenotype," signifying the similarity between the transcriptional state induced by the TF increase or decrease and the transcriptional state of a specific blood lineage cell type.

Such analyses recapitulate known biology, with *GFI1B* upregulation ^29^ and *MYB* downregulation ^44^ being crucial factors promoting erythrocyte maturation (**Figure 5A**). The downregulation of *NFE2* instead was negatively related to platelet differentiation. Analysing the correlations as inferred phenotypes suggests potential nonlinear relationships (**Figure S12B**), but these trends should be considered hypotheses that require experimental validation. In summary, this points to cellular phenotypes resulting from gradual TF dosage modulation.

**Figure 5:**
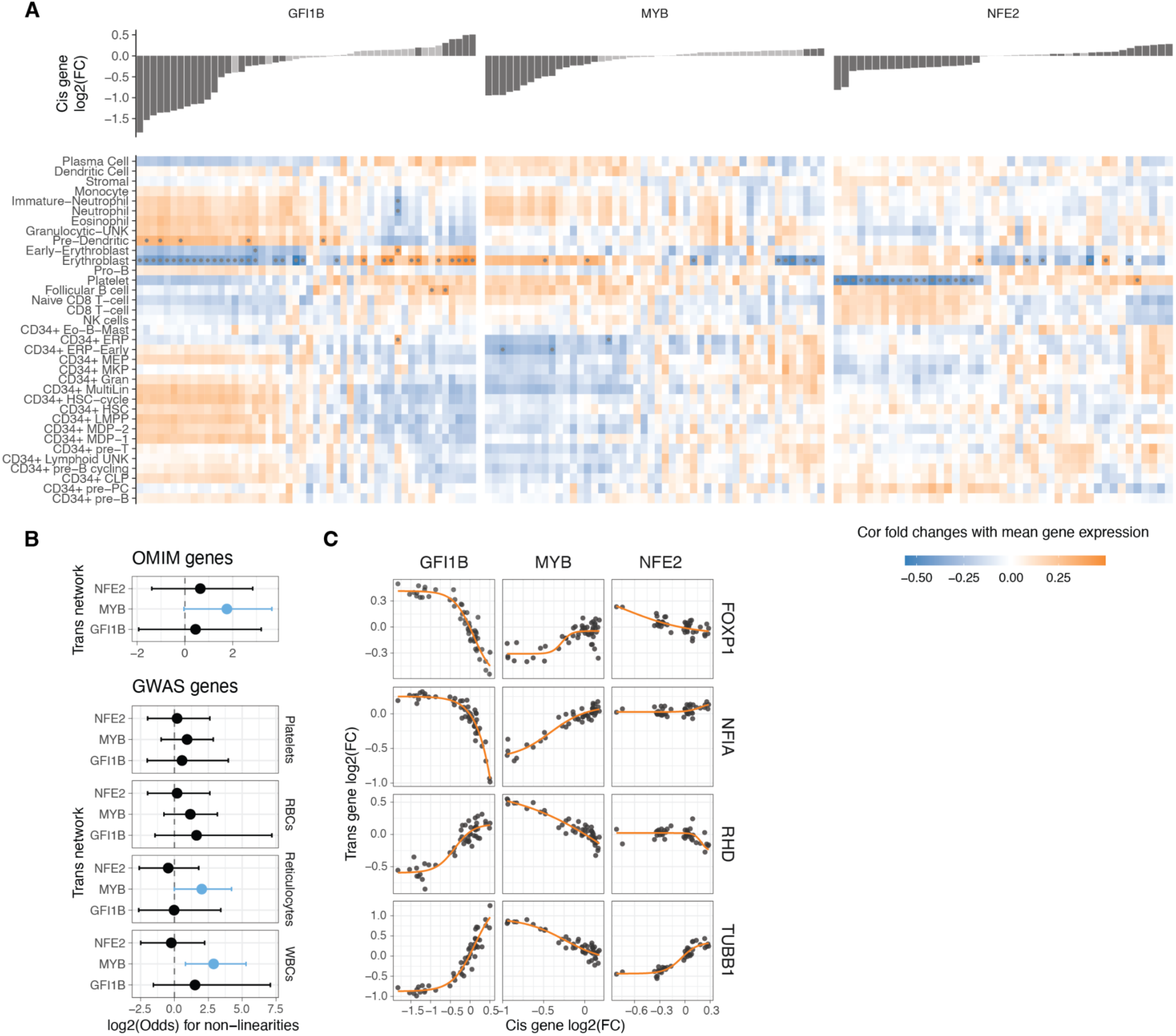
Non-linearities in TF dosage responses of complex traits and disease genes. A. Heatmap illustrating the correlation between the mean expression of cell types and the changes in expression linked to individual TF dosage perturbations. The barplot on the top panel represents cis gene dosage perturbation. Asterisks (*) denote correlations with 10% FDR. B. Enrichment log(odds) ratio of non-linear TF dosage responses (ΔAIC_linear-sigmoid_ > 0) in disease related genes (OMIM genes linked to 1 or more diseases, top panel) or in GWAS blood traits associated genes (closest expressed gene to lead GWAS variant, bottom panel). Log(odds) with Fisher’s exact test at FDR < 0.05 are highlighted in blue. C. Examples of TF dosage response curves of genes both associated with disease (OMIM) and complex traits (Blood GWAS).

Many of the analysed *trans* genes are associated with physiological traits and diseases (**Figure 4**). Understanding the nonlinear trends in the expression of these genes is of particular interest, as it helps comprehend how genes with physiological impacts may be buffered against upstream regulatory changes, and how their dosage changes as a response to upstream regulators contrasts with genetic variants that contribute to diseases and traits. Additionally, knowing the underlying dosage-to-phenotype curve of a gene can be crucial if this is considered a biomarker for identifying or treating disease. To investigate this, we analysed whether OMIM genes for rare diseases and Mendelian traits or GWAS genes for different blood cell traits (**Figure 5B**) that are part of the *trans* networks of genes affected by *GFI1B*, *MYB,* or *NFE2* perturbation are enriched for nonlinear dosage responses. As seen in **Figure 4**, the *trans* response properties of each gene are highly specific to the regulators and thus analysed in parallel for each *cis* gene network. An enrichment for nonlinear responses was observed for *MYB trans* network genes associated with disease and for blood traits related to white blood cells and reticulocytes. These enrichments are particularly interesting given that most *trans* genes that were sensitive to *MYB* dosage modulation did not respond with a nonlinear trend (**Figure S9A**).

Despite nonlinear responses not being significantly enriched among disease genes across all *trans* networks, the responses of the same *trans* gene can show very different dosage responses depending on the upstream regulator being tuned. In Figure 5C, we highlight several disease-associated genes (linked to one or more disease phenotypes ^45^. *FOXP1*, a haploinsufficient and potentially triplosensitive transcription factor implicated in intellectual disability, exhibited a strong and dose-dependent response, particularly to varying levels of *GFI1B*. A similar pattern was observed for *NFIA*, another haploinsufficient gene involved in developmental disorders.. However, it is difficult to interpret their expression response in K562 cells when their most apparent phenotypic effects likely derive from other cell types. *RHB* is the Rhesus blood type gene, where a common deletion of the gene causes the Rh-blood type in homozygous individuals, with a strong nonlinear response to *GFI1B* levels. A particularly interesting gene is *TUBB1*, part of β-tubulin, that causes autosomal dominant macrothrombocytopenia or abnormally large platelets. Here, K562 cells are a reasonable model system, being closely related precursors to megakaryocytes that produce platelets. Interestingly, *GFI1B* loss also causes a macrothrombocytopenia phenotype in mice ^46^, and in our data, *TUBB1* expression decreases quickly as a function of decreased *GFI1B* expression but then plateaus at a level that corresponds to loss of one copy of *TUBB1*. This raises the hypothesis that low *GFI1B* levels may cause macrothrombocytopenia at least partially via reducing *TUBB1* expression.

## Discussion

In this paper, we have investigated how gradual dosage modulation of transcription factors contributes to dosage-sensitive transcriptional regulation and investigated its potential phenotypic consequences. First, we set up an easily scalable and generalizable CRISPRi/CRISPRa approach with tiling sgRNAs for gradual titration of gene expression, with reagents that can be designed with data only of the TSS and easily ordered at scale. Alternative approaches that rely e.g., on targeting CREs that are often unknown, dramatic overexpression, or laborious setting up of constructs for each gene are less practical for large-scale analyses. Our approach appears best suited for expression modulation in the biologically reasonable range, and other methods would be needed for dramatic overexpression or complete silencing of the target genes. Our inability to substantially increase *MYB* expression indicates the need for further work and larger *cis* gene sets to fully understand how widespread this is and to what extent this depends on *cis*-regulatory properties versus feedback and buffering mechanisms. Nevertheless, we believe that the approach proposed here is a useful complement to the diversifying set of tools for dosage modulation for different purposes ^18–22^.

In this work, we made use of targeted transcriptome sequencing to avoid complications from the sparsity of single-cell data. While highly accurate targeted readout of the *cis* gene expression linked to each sgRNA is a core component of our approach, analysing *trans* responses could also be achieved by standard single-cell sequencing of the full transcriptome, possibly in combination with a targeted readout of transcripts of particular interest. In this study, the targeted genes were selected based on prior data of responding to *GFI1B*, *NFE2,* or *MYB* regulation and thus do not represent an unbiased or random sample of genes. An interesting future extension would be the addition of single-cell protein quantification to confirm that the detected mRNA levels correspond to protein levels, but this remains technically challenging.

Our results show that nonlinear responses to gradual up- and down-regulation of TF dosage are widespread and can be detected even without extreme overexpression or full knockout of the TF. The patterns of transcriptional responses are highly context-specific and vary between upstream regulators. Further work with larger sets of *cis* and *trans* genes, as well as direct quantification of cellular readouts, will be needed to fully characterise the patterns and mechanisms of downstream impacts on gene dosage. However, our findings indicate important directions for future research. First, the widespread nonlinearity suggests that inferring gene function from classical molecular biology approaches—such as drastic knockouts or knockdowns—may be limited, as these perturbations can produce effects that are both quantitatively and qualitatively different from the more modest changes that occur naturally. This may be particularly relevant for essential and highly dosage-sensitive genes, where applying our gradual dosage modulation framework can provide opportunities for functional characterization at perturbation levels that do not kill the cells. Secondly, we show that the effects of up- and downregulation are qualitatively and quantitatively different, which calls for increased attention to analysing both directions of effect, which also occur in natural responses to genetic variants and environmental stimuli.

Gene dosage sensitivity has typically been studied by human genetics and genomics methods ^47–49^. The experimental approach pursued in this study and the computational approaches are fundamentally different and complement each other: while human genetics is powerful for capturing the functional importance of physiological phenotypes via patterns of population variation and selective constraint, experimental approaches provide more granularity and insights into cellular mechanisms. Furthermore, while the convergence of disease effects of common and rare variants affecting the same gene is a well-known phenomenon ^6,7^, the sparsity of variants makes it difficult to properly model allelic series as a continuous dosage-to-phenotype function for individual genes. Experimental approaches can provide a powerful complement to this. Altogether, we envision that combining these perspectives into true systems genetics approaches will be a powerful way to understand how gene dosage variation contributes to human phenotypes from molecular to cellular and eventually physiological levels.

## Acknowledgments

This work was funded by NIH grants R01MH106842, R01AG057422, DP2HG010099, R01HG012790, and R01GM122924; a grant from the Knut and Alice Wallenberg Foundation to SciLifeLab for research in Data-driven Life Science, DDLS (KAW 2020.0239); funding from the European Research Council (ERC) under the European Union’s Horizon 2020 research and innovation programme (Grant agreement No. 101043238); a European Molecular Biology Organization Postdoctoral Fellowship (ALTF 345-2021) to J.D.; a Canadian Institutes of Health Research Banting Postdoctoral Fellowship and NIH/NHGRI (K99HG012792) to J.A.M.

## Author Contributions

J.D. and T.L. conceived the study. J.D. performed the experiments, with contributions from J.A.M. and M.Z.. N.S. contributed experimental resources to the study. J.D. performed the computational analyses, with contributions from M.M., S.G. and J.A.M.. J.D. and T.L. wrote the manuscript with contributions and review from all the authors.

## Data availability and materials

All code used in this study is available at https://github.com/LappalainenLab/d2n_ms. Raw sequencing data has been submitted to GEO (accession number GSE257547).

## References

1. Maurano, M. T. et al. Systematic localization of common disease-associated variation in regulatory DNA. Science 337, 1190–1195 (2012).

2. GTEx Consortium. The GTEx Consortium atlas of genetic regulatory effects across human tissues. Science 369, 1318–1330 (2020).

3. Claussnitzer, M. et al. A brief history of human disease genetics. Nature 577, 179–189 (2020).

4. Morris, J. A. et al. Discovery of target genes and pathways at GWAS loci by pooled single-cell CRISPR screens. Science 380, eadh7699 (2023).

5. Zschocke, J., Byers, P. H. & Wilkie, A. O. M. Mendelian inheritance revisited: dominance and recessiveness in medical genetics. Nat. Rev. Genet. 24, 442–463 (2023).

6. Backman, J. D. et al. Exome sequencing and analysis of 454,787 UK Biobank participants. Nature 599, 628–634 (2021).

7. Freund, M. K. et al. Phenotype-Specific Enrichment of Mendelian Disorder Genes near GWAS Regions across 62 Complex Traits. Am. J. Hum. Genet. 103, 535–552 (2018).

8. Castel, S. E. et al. Modified penetrance of coding variants by cis-regulatory variation contributes to disease risk. Nat. Genet. 50, 1327–1334 (2018).

9. Fahed, A. C. et al. Polygenic background modifies penetrance of monogenic variants for tier 1 genomic conditions. Nat. Commun. 11, 3635 (2020).

10. McCaw, Z. R. et al. An allelic-series rare-variant association test for candidate-gene discovery. Am. J. Hum. Genet. 110, 1330–1342 (2023).

11. Sanjana, N. E. Genome-scale CRISPR pooled screens. Anal. Biochem. 532, 95–99 (2017).

12. Collins, R. L., et al. A cross-disorder dosage sensitivity map of the human genome. medRxiv (2021).

13. Fehrmann, R. S. N. et al. Gene expression analysis identifies global gene dosage sensitivity in cancer. Nat. Genet. 47, 115–125 (2015).

14. Wang, T. et al. Identification and characterization of essential genes in the human genome. Science 350, 1096–1101 (2015).

15. Hart, T. et al. High-Resolution CRISPR Screens Reveal Fitness Genes and Genotype-Specific Cancer Liabilities. Cell 163, 1515–1526 (2015).

16. Cowley, G. S. et al. Parallel genome-scale loss of function screens in 216 cancer cell lines for the identification of context-specific genetic dependencies. Sci Data 1, 140035 (2014).

17. Keren, L. et al. Massively Parallel Interrogation of the Effects of Gene Expression Levels on Fitness. Cell vol. 166 1282–1294.e18 Preprint at 10.1016/j.cell.2016.07.024 (2016).

18. Jost, M. et al. Titrating gene expression using libraries of systematically attenuated CRISPR guide RNAs. Nat. Biotechnol. 38, 355–364 (2020).

19. Hawkins, J. S. et al. Mismatch-CRISPRi Reveals the Co-varying Expression-Fitness Relationships of Essential Genes in Escherichia coli and Bacillus subtilis. Cell Syst 11, 523–535.e9 (2020).

20. Noviello, G., Gjaltema, R. A. F. & Schulz, E. G. CasTuner is a degron and CRISPR/Cas-based toolkit for analog tuning of endogenous gene expression. Nat. Commun. 14, 3225 (2023).

21. Chiarella, A. M. et al. Dose-dependent activation of gene expression is achieved using CRISPR and small molecules that recruit endogenous chromatin machinery. Nat. Biotechnol. 38, 50–55 (2020).

22. Liu, W. et al. Dissecting reprogramming heterogeneity at single-cell resolution using scTF-seq. bioRxiv 2024.01.30.577921 (2024) doi:10.1101/2024.01.30.577921.

23. Lalanne, J.-B., Parker, D. J. & Li, G.-W. Spurious regulatory connections dictate the expression-fitness landscape of translation factors. Mol. Syst. Biol. 17, e10302 (2021).

24. Naqvi, S. et al. Precise modulation of transcription factor levels identifies features underlying dosage sensitivity. Nat. Genet. (2023) doi:10.1038/s41588-023-01366-2.

25. Pulice, J. L. & Meyerson, M. Dosage amplification dictates oncogenic regulation by the NKX2-1 lineage factor in lung adenocarcinoma. bioRxiv (2023) doi:10.1101/2023.10.26.563996.

26. Mostafavi, H., Spence, J. P., Naqvi, S. & Pritchard, J. K. Systematic differences in discovery of genetic effects on gene expression and complex traits. Nat. Genet. 55, 1866–1875 (2023).

27. van der Lee, R., Correard, S. & Wasserman, W. W. Deregulated Regulators: Disease-Causing cis Variants in Transcription Factor Genes. Trends Genet. 36, 523–539 (2020).

28. Ulirsch, J. C. et al. Systematic Functional Dissection of Common Genetic Variation Affecting Red Blood Cell Traits. Cell 165, 1530–1545 (2016).

29. Möröy, T., Vassen, L., Wilkes, B. & Khandanpour, C. From cytopenia to leukemia: the role of Gfi1 and Gfi1b in blood formation. Blood 126, 2561–2569 (2015).

30. Polfus, L. M. et al. Whole-Exome Sequencing Identifies Loci Associated with Blood Cell Traits and Reveals a Role for Alternative GFI1B Splice Variants in Human Hematopoiesis. Am. J. Hum. Genet. 99, 481–488 (2016).

31. Jutzi, J. S. et al. Altered NFE2 activity predisposes to leukemic transformation and myelosarcoma with AML-specific aberrations. Blood 133, 1766–1777 (2019).

32. Baker, S. J. et al. B-myb is an essential regulator of hematopoietic stem cell and myeloid progenitor cell development. Proc. Natl. Acad. Sci. U. S. A. 111, 3122–3127 (2014).

33. Doench, J. G. et al. Rational design of highly active sgRNAs for CRISPR-Cas9-mediated gene inactivation. Nat. Biotechnol. 32, 1262–1267 (2014).

34. Doench, J. G. et al. Optimized sgRNA design to maximize activity and minimize off-target effects of CRISPR-Cas9. Nat. Biotechnol. 34, 184–191 (2016).

35. McKenna, A. & Shendure, J. FlashFry: a fast and flexible tool for large-scale CRISPR target design. BMC Biol. 16, 74 (2018).

36. Sanson, K. R. et al. Optimized libraries for CRISPR-Cas9 genetic screens with multiple modalities. Nat. Commun. 9, 5416 (2018).

37. Legut, M. et al. High-Throughput Screens of PAM-Flexible Cas9 Variants for Gene Knockout and Transcriptional Modulation. Cell Rep. 30, 2859–2868.e5 (2020).

38. Luo, Y. et al. New developments on the Encyclopedia of DNA Elements (ENCODE) data portal. Nucleic Acids Res. 48, D882–D889 (2020).

39. Lupo, A. et al. KRAB-Zinc Finger Proteins: A Repressor Family Displaying Multiple Biological Functions. Curr. Genomics 14, 268–278 (2013).

40. Minaeva, M., Domingo, J., Rentzsch, P. & Lappalainen, T. Specifying cellular context of transcription factor regulons for exploring context-specific gene regulation programs. NAR Genom Bioinform. 2025 Jan 7;7(1):lqae178. doi: 10.1093/nargab/lqae178.

41. Wang, H. et al. Dynamic transcriptomes of human myeloid leukemia cells. Genomics 102, 250–256 (2013).

42. Salvadores, M., Fuster-Tormo, F. & Supek, F. Matching cell lines with cancer type and subtype of origin via mutational, epigenomic, and transcriptomic patterns. Sci Adv 6, (2020).

43. Granja, J. M. et al. Single-cell multiomic analysis identifies regulatory programs in mixed-phenotype acute leukemia. Nat. Biotechnol. 37, 1458–1465 (2019).

44. Jayapal, S. R. et al. Down-regulation of Myc is essential for terminal erythroid maturation. J. Biol. Chem. 285, 40252–40265 (2010).

45. Amberger, J. S., Bocchini, C. A., Schiettecatte, F., Scott, A. F. & Hamosh, A. OMIM.org: Online Mendelian Inheritance in Man (OMIM®), an online catalog of human genes and genetic disorders. Nucleic Acids Res. 43, D789–98 (2015).

46. Beauchemin, H. et al. Gfi1b controls integrin signaling-dependent cytoskeleton dynamics and organization in megakaryocytes. Haematologica 102, 484–497 (2017).

47. Collins, R. L. et al. A structural variation reference for medical and population genetics. Nature 581, 444–451 (2020).

48. Mohammadi, P. et al. Genetic regulatory variation in populations informs transcriptome analysis in rare disease. Science 366, 351–356 (2019).

49. Dong, D. et al. An RNA-informed dosage sensitivity map reflects the intrinsic functional nature of genes. Am. J. Hum. Genet. 110, 1509–1521 (2023).

