## Supplemental material for "Nonlinear transcriptional responses to gradual modulation of transcription factor dosage"

### Experimental methods

#### Plasmids, bacteria strains and cell lines used

Plasmids
- pCC_05: Lentiviral Puromycin CRISPRa dCas9-VPR system (Addgene 139090)
- pGC02: Lentiviral Blasticydin plasmid with CRISPRi KRAB-dCas9-MeCP2 system (Addgene 170068)
- pJDE003: Lentiviral Blasticydin CRISPRa dCas9-VPR system (this study)
- pGC03: Lentiviral Puromycin sgRNA library cloning vector (Addgene 170069)
- pMD2G: Lentiviral envelope plasmid (Addgene 12259)
- psPAX2: Lentiviral packaging plasmid (Addgene 12260)
-

Bacteria *E.coli* strains
- 5-alpha competent cells (NEB C2987H)
- One Shot Stbl3 competent cells (Invitrogen 1934665)
- Endura electrocompetent cells (Lucigen 60242-2)

Cell lines:
- HEK293FT (Thermo Fisher Scientific R70007); cells were maintained at 37℃ and 5% $CO_2$ in high glucose DMEM (Cytiva SH30022.01) supplemented with 10% Serum Plus II (Sigma-Aldrich 14009C)
- K562 (ATCC, CCL243);
  Cells were maintained at 37℃ and 5% $CO_2$; HEK293FT were cultured in high glucose DMEM (Cytiva SH30022.01) supplemented with 10% Serum Plus II (Sigma-Aldrich 14009C); K562 were cultured in IMDM, GlutaMAX (Thermo Scientific 31980097) supplemented with 10% Serum Plus II.

#### CRISPRa vector construction

To construct the vector harbouring the CRISPRa system (pJDE003), the CRISPRi (KRAB-dCas9-MeCP2) gene fusion of pGC02 [1] was replaced with dCas9-VPR cassette, which was PCR amplified (Q5 High-Fidelity 2X Master Mix, NEB M0492L) from the plasmid pCC_05 [2] with primers oJDE005 and oJDE006 following instructions from manufacturer. pGC02 was digested with XbaI-FD and BamHI-FD (Thermo Fisher FD0685 and FD0054) and sequentially dephosphorylated with FastAP (Thermo Fisher EF0651) following the manufacturer's recommendations. The digested pGC02 vector and the PCR insert with the CRISPRa system (previously treated with a 15 min DpnI enzyme incubation, Thermo Fisher FD1704) were assembled by Gibson assembly using a 2:1 insert:vector ratio with Gibson Assembly Master Mix (NEB E2611S). Assemblies were transformed into NEB 5-alpha *E.coli* competent cells and single colonies were picked and sequence validated by Sanger sequencing. Frozen stock of the correct construct cells were regrown for plasmid Maxiprep extraction (QIAGEN 12362) for subsequent virus production.

### CRISPRa K562 cell line construction and functional validation

Lentivirus was produced by polyethylenimine linear MW 25000 (Polysciences 23966) transfection of HEK293FT cells with the transfer plasmid containing a Cas9-VPR effector, packaging plasmid psPAX2 (Addgene 12260) and envelope plasmid pMD2.G (Addgene 12259). After 72 h post-transfection, cell media containing lentiviral particles was harvested and filtered through 0.45 µm filter Steriflip-HV (Millipore SE1M003M00). One volume of Lentivirus Precipitation Solution (Alstem VC100) was added to the collected lentivirus, mixed and stored overnight at 4C. The mix was centrifuged for 30 min at 1,500g, and the pellet of lentiviral particles were resuspended in 1/10th of the original volume of DMEM media. Lentivirus vials were frozen at -80C and later thawed for transduction.

To construct the monoclonal K562 cell line with the CRISPRa system, the dCas9-VPR lentivirus was transduced into one million K562 cells using 100 µl of 10X concentrated lentivirus in a total volume of 1 ml (high MOI). After 24 hours, the media was replaced with fresh IMDM, and 48 hours after transduction, blasticidin (A.G. Scientific B-1247) was added to a final concentration of 10 µg/µl for 16 days. Monoclonal cell lines were sorted by FACS (Sony Cell Sorter SH800) into a 96-well plate. The presence of dCAS9 protein in several growing clones was confirmed by western blot (Primary antibody: Purified anti-CRISPR CAS9 antibody; BioLegend 844302. Secondary antibody: LI-COR 925-32212) and protein levels were normalised to GAPDH (Primary antibody: GAPDH (14C10) Rabbit; Cell Signalling Technology 2118S. Secondary antibody: LI-COR 925-68073).

To select the final monoclonal CRISPa cell line, the three clones with the highest protein expression in the western blot were subjected to functional validation to test for activation activity. Lentiviral guides designed from [2] targeting CD4 (Anti-CD4 Mouse Monoclonal Antibody (FITC), BioLegend, 300505), which is lowly expressed in K562, CD19 (Anti-CD19 Mouse Monoclonal Antibody (APC),BioLegend, 302211) with null expression, and CD45 (PE anti-human CD45, BioLegend, 368509) with intermediate expression, were independently transduced into all three monoclonals, and after puromycin selection, the expression of this markers was screened by FACS at day 4 and at day 10 or 11 after transduction. The clone with the strongest and most consistent activity was selected.

### Gene selection for targeted sequencing and design of probe custom panel

The four selected dosage genes were GFI1B, NFE2, MYB, and TET2. GFI1B and NFE2 were chosen due to their reported trans effects following the inhibition of their cis-CREs [3]. MYB was selected for being downstream of the GFI1B network, and TET2 was selected as an unrelated gene to those transcriptional networks. Both MYB and TET2 qualified as oncogene and tumour suppressor functions in K562 (https://depmap.org/portal/), making them ideal choices to determine the impact of growth effects on the experiment.

The additional 88 genes captured by targeted sequencing were selected based on the significant trans effects of GFI1B and NFE2 inhibition from [3] including two control genes, GAPDH, and LHX3 genes, one highly constantly expressed “housekeeping gene” and the other with no reported expression in K562 cell lines, respectively. The remaining 86 genes were selected based on Morris et al. to include 1) 29 genes that overlapped between the NFE2 and GFI1B network, 2) 47 trans genes for GFI1B, 10) trans genes for NFE2. The number of trans genes selected from each unique network was proportionally chosen, given the size of each trans network, and oversampling TFs and TF targets as defined in [4], as well as maintaining the proportional co-expression cluster structure as defined in [3] (**Figure S1a,b**). Additional filters in the selection included a minimum expression of 0.1 mean UMI/cell and the lack of alternative 5' splice isoforms and a unique Ensembl ID.

The 10X Probe Full Custom Panel design tool was used to design the targeted gene expression probe library. A total of 687 probes (~15%) were discarded because they covered transcript regions with a median coverage per base of < 3 reads/bp (for medium and highly expressed genes) or <1 reads/bp (for lowly expressed genes). All probes for LHX3 (0 median reads/bp) were retained. In addition, 93 probes covering the entire transcript sequence of the dCas9-VPR and KRAB-dCas9-MeCP2 transcript were included, resulting in a final total of 4,405 probes. The xGen™ Custom Hybridization Capture Panel of biotinylated oligos was ordered and synthesised at IDT.

### Gene dosage sgRNA library design and cloning

The sgRNA library contained a total of 96 guides (51 tiling, 8 TSS, 20 attenuated, 12 enhancer and 5 non-targeting controls). All guides were designed to not contain the U6 terminator sequence, repeats of five or more consecutive G, C or As, as well as not falling in the genomic region where K562 cell line has alternative alleles compared to the human genome reference (Hg38). All guides were scored with FlashFry [5] to obtain off-target and on-target activity scores that allowed the selection of the best scoring guides. Tiling guides were designed to target different regions of the promoter, TSS and beginning of the gene body of each dosage gene, spanning a total average distance of 1400 bp (TSS in the centre), each being on average distant from one another of 110 bp. The sequences of the two TSS guides were obtained from [6]. The sgRNAs targeting enhancers were picked from previously reported work that showed a CRISPR-based evidence of enhancer activity (GFI1B [1], NFE2 [7], MYB [8]). The five attenuated guides for each gene were manually designed following the rules described in [9] to span a range of activities, including a single point mutation on the best scoring guide that targeted the TSS.

Overhangs with homology regions to the pGC03 plasmid (18bp downstream and 22 bp upstream) were added to the sgRNA sequence to be able to directly clone the ssDNA oligos into the plasmid. The 96 sgRNAs were ordered in IDT as single stranded DNA oligos (total 60bp) in a 96 well-plate to 100 pmol scale. The oligos were pooled at equimolar concentration and diluted to a final concentration of 0.2 uM. The library was cloned into the BsmBI digested plasmid pGC03 using 10 reactions of the NEBuilder HiFi DNA Assembly kit following the manufacturer’s instructions. All the reactions were pooled and the DNA precipitated using Isopropanol, GlycoBlue (Thermo Scientific AM9515) and 50mM NaCl for 15 min at RT. Following two washes with Ethanol 70%, the assembly was resuspended with 15ul of 0.2X TE.

To transform the library into E.coli, 1ul of the assembly was mixed with 25 uL of Endura cells under manufacturer’s electroporation conditions, then plated onto 245 x 245 mm square LB

100 ug/ml Carbenicillin plates. The plates were grown ON, and >2.5e5 transformants were obtained, ensuring the complexity of the library was maintained at >1000 cells per unique sgRNA. All colonies were collected and subjected to maxiprep using the Maxi Fast-Ion Plasmid Kit, Endotoxin Free kit (IBI Scientific IB47123). The representation of the library was assessed through MiSeq shallow sequencing (Illumina).

#### sgRNA lentiviral library production and cell culture assay

The lentiviral library was produced by transfecting ~80 million HEK293FT cells with a transfer plasmid containing the 96 sgRNA library, along with the packaging plasmid psPAX2 and envelope plasmid pMD2.G, using polyethylenimine linear MW 25000. The supernatant media was replaced with fresh D10 10% BSA six hours after transfection, and the virus was collected and filtered through 0.45 μm filters after 48 hours. The lentiviral library was then concentrated 2X using the Precipitation Lentiviral Solution, aliquoted, and stored at -80°C for subsequent transduction.

Both CRISPRi and CRISPRa K562 cell lines were independently transduced with different titers of the lentiviral sgRNA library at a low MOI (one sgRNA per cell). Twenty hours post-transduction, the cell media was replaced with fresh IMDM 10% Serum Plus II Blasticidin 5 μg/mL, and four hours later, Puromycin (Invivogen ant-pr-1) was added at a final concentration of 2 μg/mL to select for cells with sgRNA integration. The transduction batch with an infection rate of ~10% was selected, and cells were sorted to near purity using FACS to remove dead cells. Cells were maintained at >90% survival and a maximum confluency of 700,000 cells/mL. On day 8 post-transduction, the cells were collected and prepared for cell hashing.

#### Multimodal single-cell experiment and targeted sequencing

Cell hashing was performed as previously described using four hashtag-derived oligonucleotides (HTOs) in a hyperconjugation protocol [10]. Each transduced cell line was split into four batches of 500,000 cells, resulting in a total of 8 different hashes. After incubation and washes, all 8 hashed batches were pooled together and run in two reaction lanes of the 10X Chromium Next GEM Single Cell 5’ Reagent Kit v2 (single indexing, PN-1000265 and PN-1000190). The manufacturer's protocol was followed with modifications stipulated in the ECCITE-seq protocol [11]. For each GEM reaction, 42,000 cells from the hash pool were used to obtain approximately 21,000 total cells, including "multiplets" (multiple cells per droplet counts). Gene expression (cDNA), hashtags (HTOs), and guide RNA (Guide-derived oligos, GDOs) libraries were constructed following the 10x Genomics and ECCITE-seq protocols (https://cite-seq.com/eccite-seq/) with minimal modifications. Specifically, the antibody pool protein tag library steps were ignored, and a custom-designed probe library was used to enrich the cDNA for the genes of interest in the 10X Targeted Gene Expression protocol (10X PN-1000248). The resulting libraries were sequenced using an Illumina Nextseq 500/550 Mid-Output v2.5 Kit (150 cycles). The targeted enrichment of the dCas9 transcripts was performed separately using an independent probe library and was sequenced together with additional HTO libraries using the Illumina Miseq Reagent Kit v3 (150 cycles).

### Computational and statistical analyses

#### From fastqs to QCed and demultiplexed UMI normalised matrices

FastQC was used to demultiplex the different samples of the three different modalities from the different 10X chip lanes, which each was processed independently. For the cDNA modality, the UMI count matrix was obtained using Cellranger *count*, including the *targeted-panel* argument to get the additional filtered matrices and summary statistics. Cells with less than 500 UMIs per cell or less than 50 genes with at least 1 UMI per cell were discarded. The top 1% cells containing the highest UMI content were also discarded. The expression of all genes across 10X lanes were extremely reproducible (Pearson r = 0.999), showing a ~5-fold UMI count increase in contrast to the non-targeted transcriptome (**Figure S1c**).

For the GDO modality (sgRNAs), Cellranger count was also used using the CRISPR Guide Capture Analysis mode, which uses a Gaussian mixture model to call sgRNA per cell. The cells containing more than one sgRNA were discarded.

To classify each cell into their corresponding CRISPR system of origin (CRISPRi or CRISPRa), both the HTO modality (protein hashes) and the expression of the dCas9 targeted transcript was used. Protein hashes were called using Alevin salmon [12], and the resulting HTO UMI matrix was mixed in with the cDNA matrix containing the expression of the CRISPRi and CRISPRa genes. This matrix was normalised and scaled using Seurat v4 [13] and used to generate a UMAP based on the expression of protein hashes and the dCas9 transcripts expression. Clusters were identified and manually assigned to an HTO category given the expression pattern of each cluster. Finally, the 5% cells classified as CRISPRi that had the lowest expression of CRISPRi transcript were discarded, as well as those 5% of CRISPRa classified cells that had the highest CRISPRi transcript expression. In total, 20,001 (10,647 CRISPRi and 9,354 CRISPRa) cells passed all filters and were used for subsequent analyses.

Once each single cell was classified into a unique sgRNA perturbation and to a cell line of origin, the cDNA UMI matrices of the two 10X lanes were merged and afterwards normalised using a the log1p normalisation method of Seurat's NormalizeData (Seurat version 4.3). On average, each unique sgRNA perturbation was measured in 81 and 86 cells for the CRISPRa and CRISPRi, respectively **(Figure S1d**)

#### Expression fold-change calculation and non-target sgRNA filtering

As estimates of changes in expression, we used a pseudo-bulk differential analyses approach. To get rid of the batch effects deriving from each cell line (CRISPRa vs. CRISPRa) (**Figure S3a**), for each unique perturbation we calculated log2 fold-change of the of the expression a gene against the expression of that gene in the population of cells harbouring the NTC sgRNAs of that particular cell line.

Before running the differential analyses on all targeted genes, across all unique CRISPR perturbations we identified those NTC sgRNAs that had potential unexpected off target activity and thus could not be used as negative controls. For all possible unique NTC sgRNA pairs we run the above differential expression analysis on all 92 targeted genes. We discarded NTC sgRNAs that showed more than one DE gene (FDR 0.05) in more than one pairwise comparison, and the differential genes showed consistent patterns of change in expression. For this reason, cells harbouring sgRNA NTC_2 on the CRISPRa modality were discarded, as this particular perturbation showed consistent undesired activation of PPP1R14A and CTCFL genes. Additionally, we ran Sceptre [14] using the resulting group of control cells to validate that our samples were calibrated correctly (**Figure S1e**).

Once those potential outlier NTCs were discarded, the log2FC of each targeted gene in each unique sgRNA and cell line condition was calculated. We used Seurat's FindMarkers() function, which computes the log fold change as the difference between the average normalized gene expression in each group on the natural log scale:

Logfc = log_e(mean(expression in group 1)) - log_e(mean(expression in group 2))

This is calculated in pseudobulk where cells with the same sgRNA are grouped together and the mean expression is compared to the mean expression of cells harboring NTC guides. To calculate per-gene differential expression p-value between the two cell groups (cells with sgRNA vs cells with NTC), Wilcoxon Rank-Sum test was used. Adjusted FDR p-values were calculated across all tests to later on call significance on DE genes. The obtained fold-changes and FDRs were used for all subsequent analyses.

#### Linear, loess and sigmoidal model fitting

To identify the best predictive model of each cis-gene dosage to trans-gene fold-change, we fitted three types of models to the data: linear (using the R *lm* function), a four parameter sigmoid (using the *drm(fct = L.4())* function from the R dcr package) and a LOESS fit (R *loess* function). To evaluate and compare the goodness of fit of the linear vs. the sigmoid model taking into account overfitting, we calculated the Akaike information criterion (AIC) using the *AIC* function from the R stats package.

To obtain an accurate prediction of each trans gene expression given TF dosage and avoid overfitting, a 10-fold cross validation scheme was followed by fitting the sigmoid model individually to each curve. The data was randomly split in 10 groups, where 90% of the data was used for training and the remaining 10% for testing. To obtain the values of each individual sigmoid fit for each dosage and trans gene response, the average and standard deviation of each parameter value was calculated across the 10 trained models.

Those trans genes with a slope significantly different from 0 (FDR adjusted p-value of a z-test across the 10 fold-CV parameter outputs) and with a min-to-max range significantly higher than 0.05 (FDR adjusted p-value of a z-test across the difference between the min and max asymptotes parameters in the 10 fold-CV), were classified as as "responsive" genes. The remaining genes were classified as “unresponsive”. The top 5% trans genes of the GFI1B trans network with the largest ∆RMSE between the LOESS fit and the sigmoid fit ($RMSE_{Sigmoid}$ - $RMSE_{LOESS}$) were classified as non-monotonic and the curve trend manually validated. For the five trans genes classified to have a non-monotonic gene expression response, their predicted expression upon TF dosage change was calculated using the LOESS model instead of the sigmoid one.

#### Gene-specific properties

Diverse gene annotations and properties were collected to compare with the different *trans* genes response properties (related to Figure 4). Quantitative annotations included the gene biotype(Ensembl [15]), Housekeeping genes [16], transcription factors [4], genes associated with at least one disease (OMIM [17]) and genes associated with blood-related complex traits (obtained from [3]).

Quantitative features included the probability of being loss-of-function intolerant scores (pLI) [18] and synonymous and missense Z scores (mis z) [18,19], which were obtained from the GnomAD database. Haploinsufficiency probability scores were obtained from [20]. To obtain the number of ChIP-Seq peaks of a *cis* gene within the promoter region of trans-genes (n peak [cis gene]), we utilised the regulon generated by Minaeva et al. 2024 [21]. This regulon was created by mapping transcription factor peaks to transcription start sites (TSS) of the 50% expressed isoforms for each gene in K562 cells, with subsequent application of a ±1 Kb proximity filter. Mean expression of genes from bone marrow cell types were obtained from Hay et al. 2018 [22] and averaged across donors. The number of protein-protein interactions of each gene within the entire human proteome (Num PPIs1) was obtained from the STING database [23].

To test significant differences between groups of genes (qualitative features), the Wilcoxon rank test was used. For quantitative features, Pearson correlation between parameters from the sigmoid model with quantitative gene metrics was used. Non-responsive and non-monotonic genes in each trans network were excluded.

#### Code and data accessibility

All code used in this study is available at https://github.com/LappalainenLab/d2n_ms. Raw sequencing data has been submitted to GEO (accession number GSE257547).

1. Morris, J. A. *et al.* Discovery of target genes and pathways of blood trait loci using pooled CRISPR screens and single cell RNA sequencing. Preprint at https://doi.org/10.1101/2021.04.07.438882.

2. Legut, M. *et al.* High-Throughput Screens of PAM-Flexible Cas9 Variants for Gene

Knockout and Transcriptional Modulation. *Cell Rep.* **30**, 2859–2868.e5 (2020).

3. Morris, J. A. *et al.* Discovery of target genes and pathways at GWAS loci by pooled single-cell CRISPR screens. *Science* **380**, eadh7699 (2023).

4. Garcia-Alonso, L., Holland, C. H., Ibrahim, M. M., Turei, D. & Saez-Rodriguez, J. Benchmark and integration of resources for the estimation of human transcription factor activities. *Genome Res.* **29**, 1363–1375 (2019).

5. McKenna, A. & Shendure, J. FlashFry: a fast and flexible tool for large-scale CRISPR target design. *BMC Biol.* **16**, 74 (2018).

6. Replogle, J. M. *et al.* Mapping information-rich genotype-phenotype landscapes with genome-scale Perturb-seq. *Cell* **185**, 2559–2575.e28 (2022).

7. Xie, S., Armendariz, D., Zhou, P., Duan, J. & Hon, G. C. Global Analysis of Enhancer Targets Reveals Convergent Enhancer-Driven Regulatory Modules. *Cell Rep.* **29**, 2570–2578.e5 (2019).

8. Li, M. *et al.* Regulation of MYB by distal enhancer elements in human myeloid leukemia. *Cell Death Dis.* **12**, 223 (2021).

9. Jost, M. *et al.* Titrating gene expression using libraries of systematically attenuated CRISPR guide RNAs. *Nat. Biotechnol.* **38**, 355–364 (2020).

10. Stoeckius, M. *et al.* Simultaneous epitope and transcriptome measurement in single cells. *Nat. Methods* **14**, 865–868 (2017).

11. Mimitou, E. P. *et al.* Multiplexed detection of proteins, transcriptomes, clonotypes and CRISPR perturbations in single cells. *Nat. Methods* **16**, 409–412 (2019).

12. Srivastava, A., Malik, L., Sarkar, H. & Patro, R. A Bayesian framework for inter-cellular information sharing improves dscRNA-seq quantification. *Bioinformatics* **36**, i292–i299 (2020).

13. Hao, Y. *et al.* Integrated analysis of multimodal single-cell data. *Cell* **184**, 3573–3587.e29 (2021).

14. Barry, T., Wang, X., Morris, J. A., Roeder, K. & Katsevich, E. SCEPTRE improves calibration and sensitivity in single-cell CRISPR screen analysis. *Genome*

*Biol.* **22**, 344 (2021).

15. Martin, F. J. *et al.* Ensembl 2023. *Nucleic Acids Res.* **51**, D933–D941 (2023).

16. Eisenberg, E. & Levanon, E. Y. Human housekeeping genes, revisited. *Trends Genet.* **29**, 569–574 (2013).

17. Amberger, J. S., Bocchini, C. A., Schiettecatte, F., Scott, A. F. & Hamosh, A. OMIM.org: Online Mendelian Inheritance in Man (OMIM®), an online catalog of human genes and genetic disorders. *Nucleic Acids Res.* **43**, D789–98 (2015).

18. Lek, M. *et al.* Analysis of protein-coding genetic variation in 60,706 humans. *Nature* **536**, 285–291 (2016).

19. Samocha, K. E. *et al.* A framework for the interpretation of de novo mutation in human disease. *Nat. Genet.* **46**, 944–950 (2014).

20. Collins, R. L. *et al.* A cross-disorder dosage sensitivity map of the human genome. *Cell* **185**, 3041–3055.e25 (2022).

21. Minaeva, M., Domingo, J., Rentzsch, P. & Lappalainen, T. Specifying cellular context of transcription factor regulons for exploring context-specific gene regulation programs. *bioRxiv* 2023.12.31.573765 (2024) doi:10.1101/2023.12.31.573765.

22. Hay, S. B., Ferchen, K., Chetal, K., Grimes, H. L. & Salomonis, N. The Human Cell Atlas bone marrow single-cell interactive web portal. *Exp. Hematol.* **68**, 51–61 (2018).

23. Szklarczyk, D. *et al.* STRING v10: protein-protein interaction networks, integrated over the tree of life. *Nucleic Acids Res.* **43**, D447–52 (2015).

### Supplementary Figures

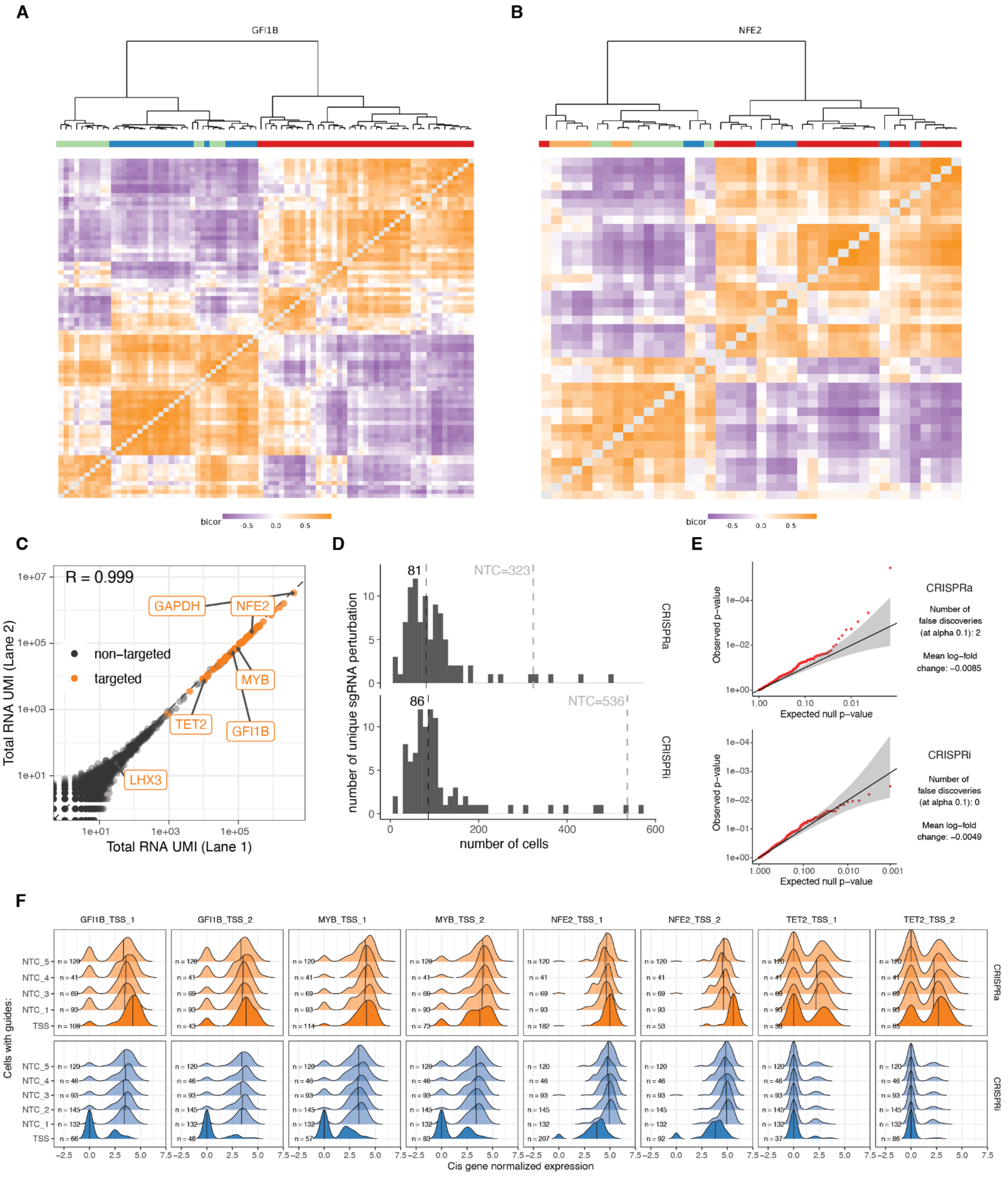

**Figure S1: Experimental design and data processing from UMIs to expression fold-change, related to Figure 1 and STAR methods.**

A. Co-expression matrix of the 76 selected GFI1B trans genes based on K562 data from [1]. Three clusters from the selected targeted panel show similar co-expression architecture than the original clusters identified using the entire GFI1B trans-network (original clusters A in blue, B in green and C in red).

B. Same as (A) for the 39 NFE2 trans genes (original clusters A in green, B in orange, C in blue and D in red).
C. Correlation between total UMI counts per gene between 10X chip lanes. Targeted panel genes are shown in orange and highlighted names correspond to dosage genes (NFE2, MYB, GFI1B and TET2) and low/high expression controls (LHX3 and GAPDH).
D. The number of singlet cells carrying each sgRNA in the two different CRISPR cell lines. NTC = non-targeting controls.
E. Q-Q plots from Sceptre calibration test.
F. Distribution of normalised UMI expression of the cis gene labelled on top for cells with sgRNAs targeting their TSS or harbouring NTC sgRNAs.

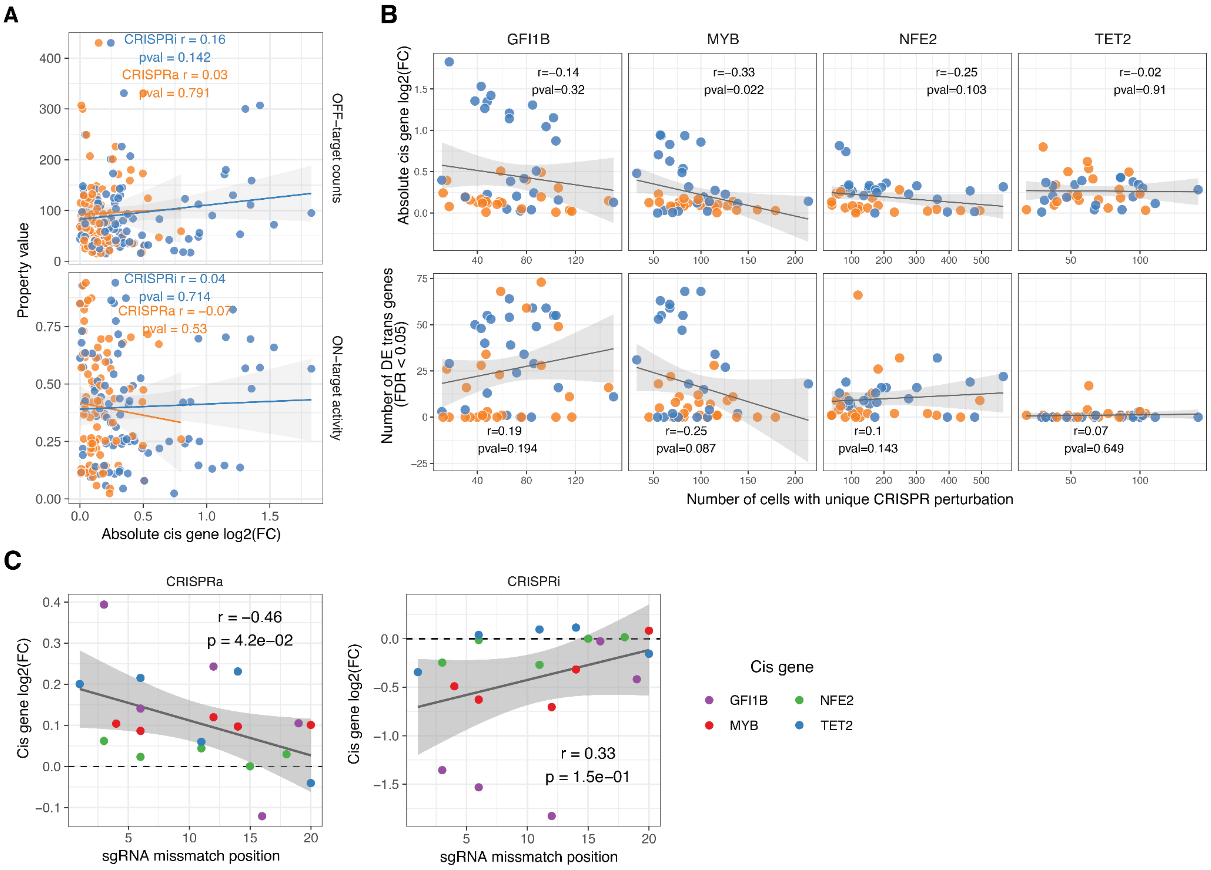


**Figure S2: Biochemical and activity properties of different types of sgRNAs**

A. Relationship between off-target and on-target activity of sgRNAs and the change in expression of their target cis gene.
B. Relationship between the number of cells that covered each sgRNA perturbation with the absolute fold change of the cis gene (top) or the number of differentially expressed trans genes due to the cis gene perturbation (bottom).
C. Relationship between the location of the mismatch mutation of attenuated sgRNAs (position 1 being farthest away from PAM motif location) and their effect on the cis gene expression.

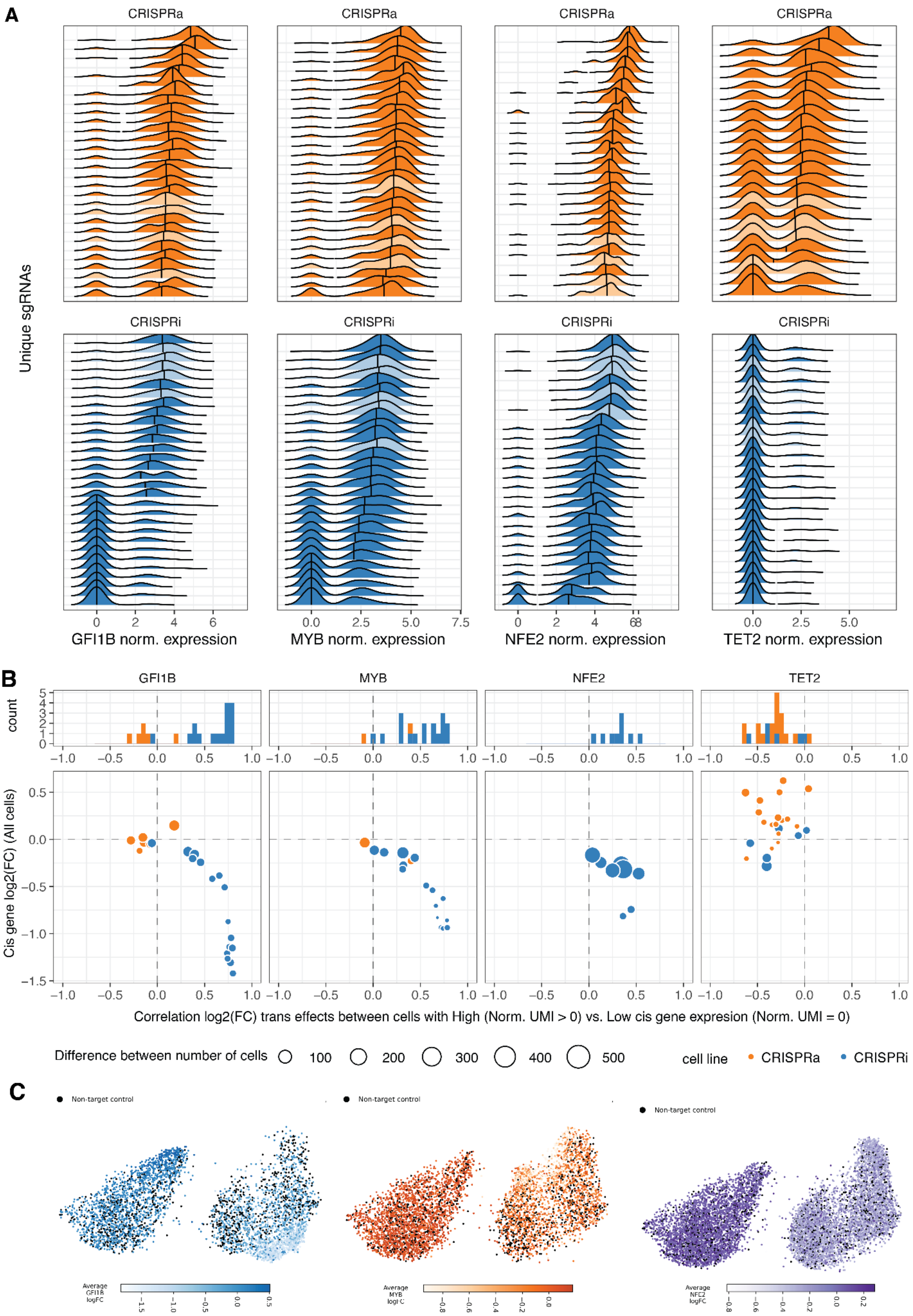


D.

**Figure S3: Gradual effects of the sgRNAs**

A. Distribution of the normalised cis gene UMIs in single cells, grouped by their unique sgRNAs, ranked top to bottom by mean normalised expression. Transparent distributions correspond to non-targeting controls.
B. Distribution of the correlation in trans gene expression fold-changes when splitting the same sgRNA cells into 0 UMI or >0 UMI for the cis gene (top panel). Comparison of the strength of these correlations with the effect of that sgRNA on the cis gene (bottom panel). Size of dots indicate the difference in the size of the 0 UMI or >0 UMI cell groups.
C. UMAPs of the cells with GFI1B, MYB, and NFE2 guides together with non-targeting guides. The left and right clusters in each figure represent CRISPRa and CRISPRi cells, respectively. The cells are coloured by the median fold change associated to their sgRNA.

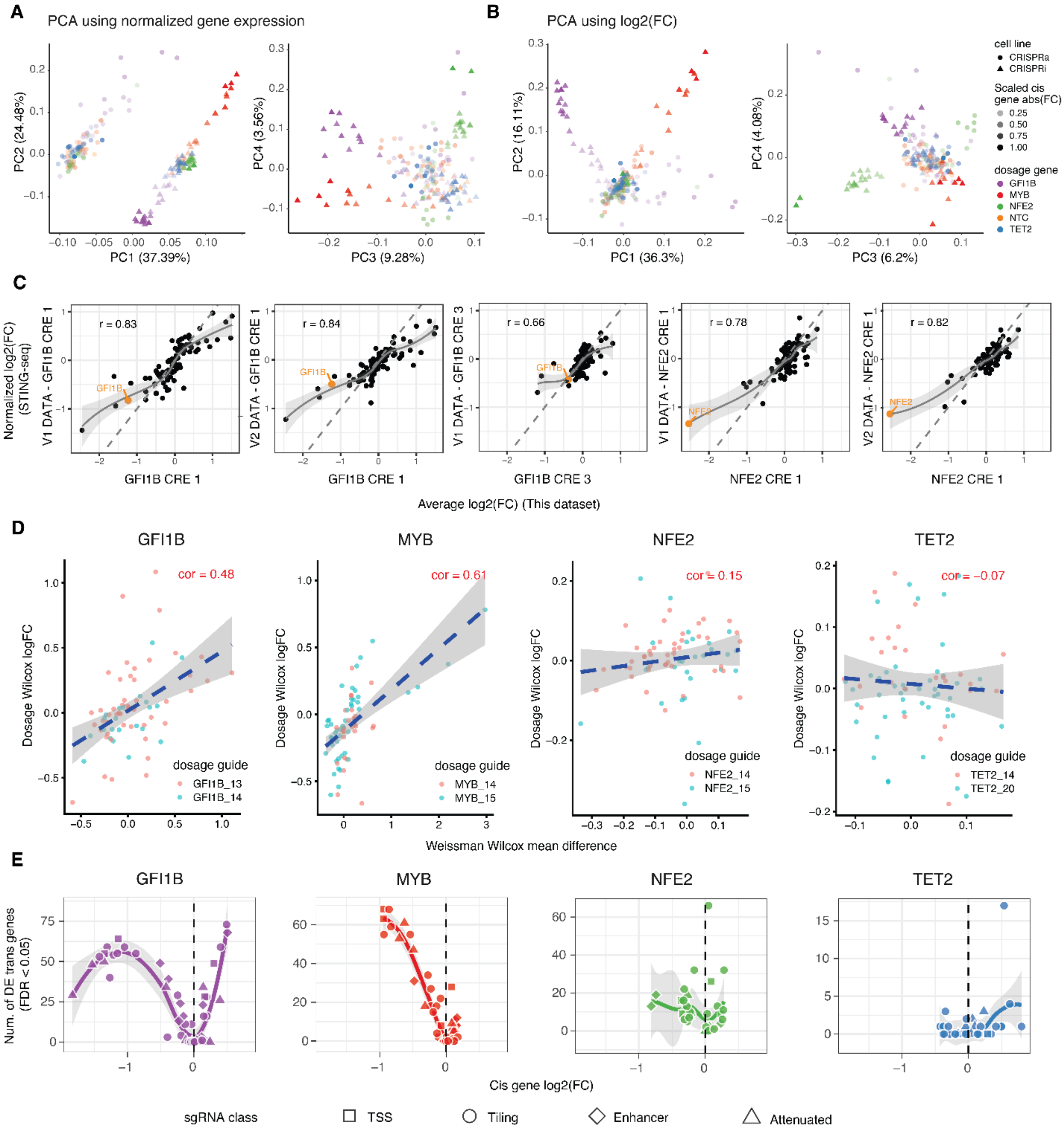

**Figure S4: Global view of trans effects and their replication**

A. PCA of mean UMI normalised expression (not relative to each cell line of origin) for all genes across unique sgRNA perturbations.

B. Same as A but using relative expression fold-change when normalising by the CRISPR cell line of origin.

C. Replication of trans-effects of CRISPRi of CREs for GFI1B and NFE2, targeted both in this study (x-axis) and in Morris et al. (y-axis). GFI1B CRE 1 and NFE2 CRE 1 were targeted in Morris et al. data batches V1 and V2, and the effects are shown here for both separately.

D. Replication of trans-effects from TSS silencing in this study and in Replogle et al., analysing guides from this study that target transcription start sites, but the guides do not fully match the exact guides used in Replogle et al. The effect size in Replogle et al. is quantified using their metric of Wilcox mean difference. The dashed line represents a linear regression line between the x and y variables.

E. Number of differentially expressed trans genes relative to the cis gene dosage perturbation.

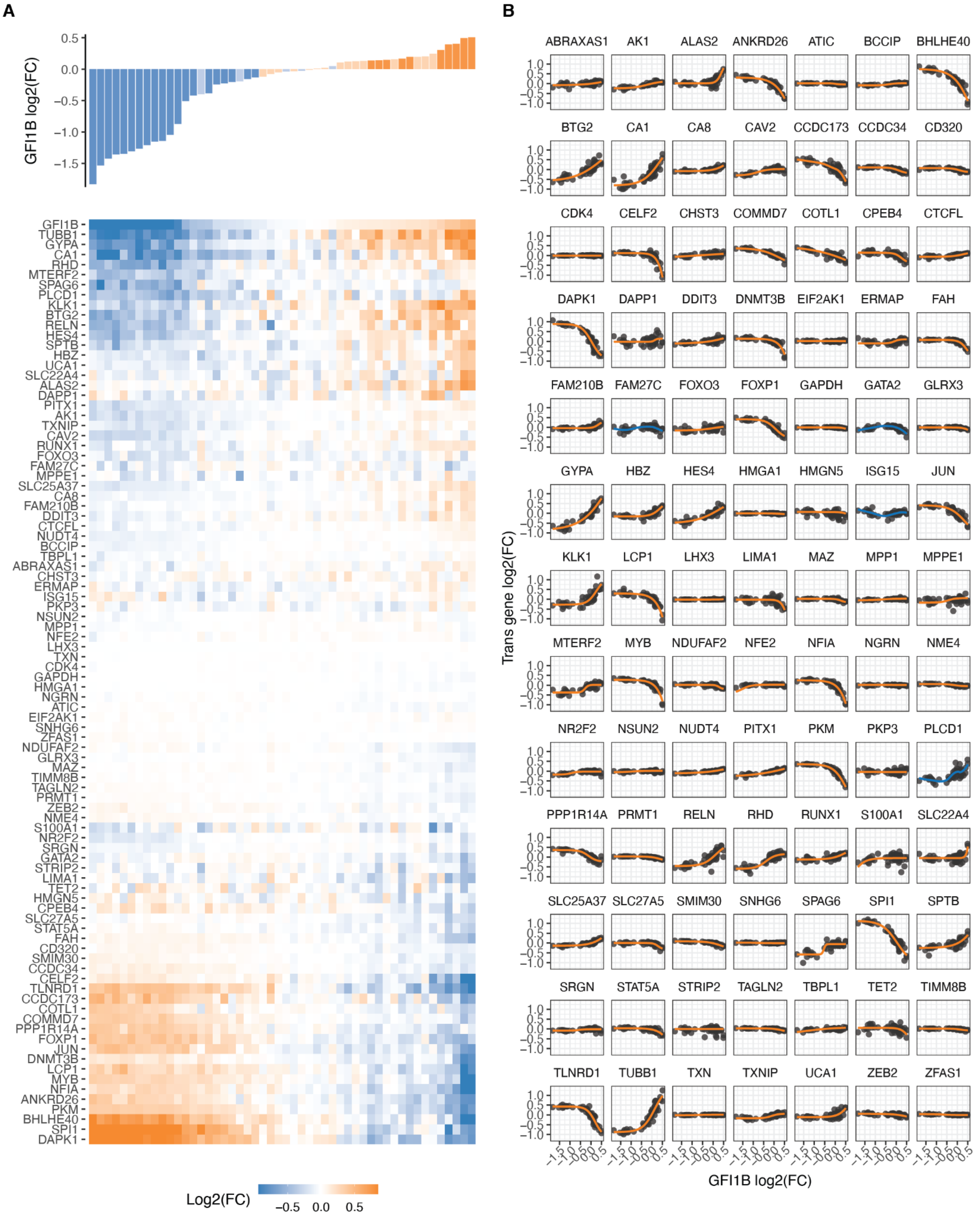


**Figure S5: Trans gene responses to GFI1B dosage modulation**

A. Changes in relative expression of all trans genes (heatmap) in response to GFI1B expression (top barplot) upon each distinct targeted sgRNA perturbation. The rows of the heatmap (trans genes) are hierarchically clustered based on their expression fold change linked to alterations in GFI1B dosage.

B. Dosage response curves are plotted for each trans gene against changes in GFI1B expression. The orange line represents the sigmoid model fit, and the blue line represents a loess curve.

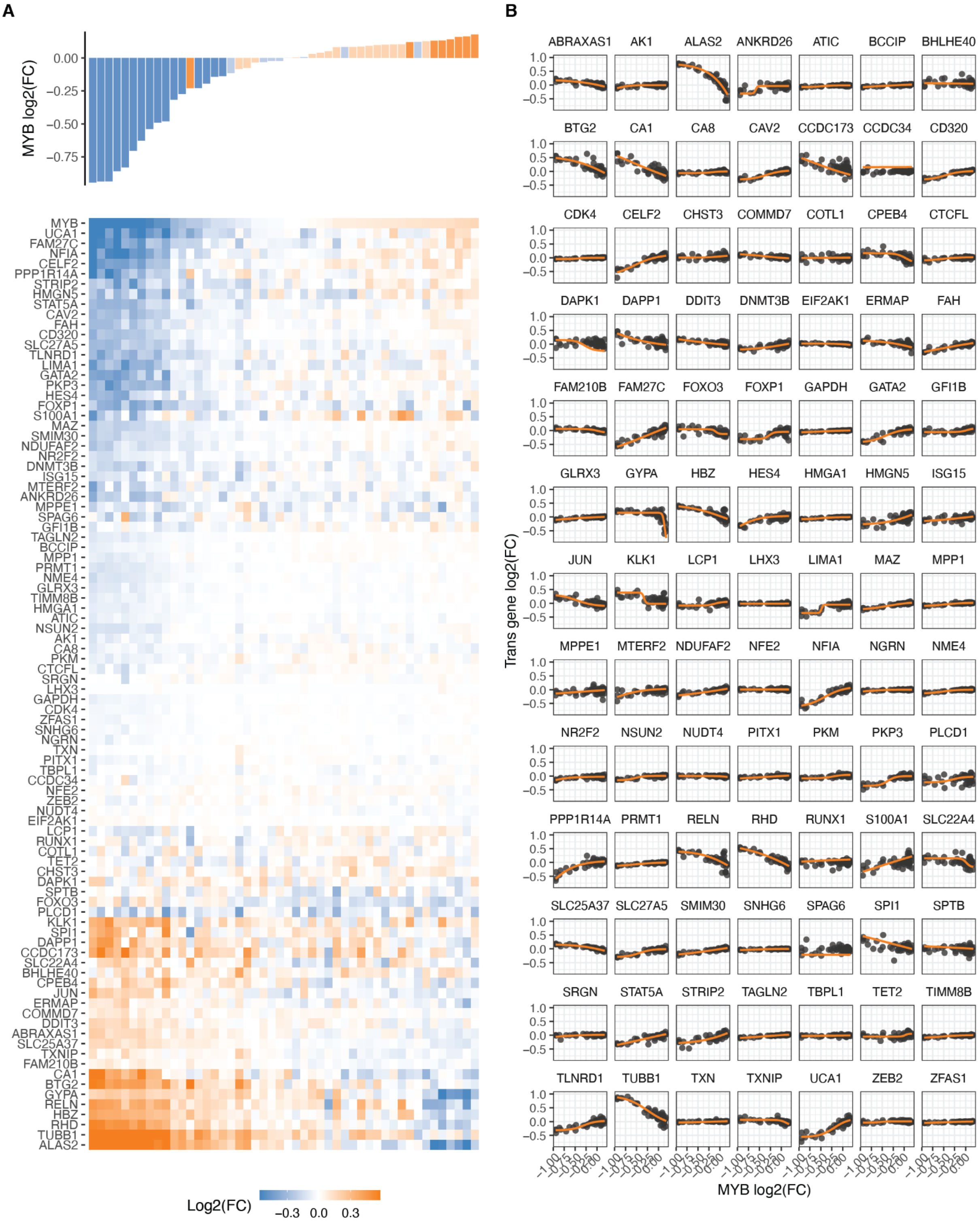


**Figure S6: Trans gene responses to MYB dosage modulation**

A. Changes in relative expression of all trans genes (bottom heatmap) in response to MYB expression (top barplot) upon each distinct targeted GFI1B sgRNA perturbation. The rows of the heatmap (trans genes) are hierarchically clustered based on their expression fold change linked to alterations in MYB dosage.
B. Dosage response curves are plotted for each trans gene against changes in MYB expression. The orange line represents the sigmoid model fit.

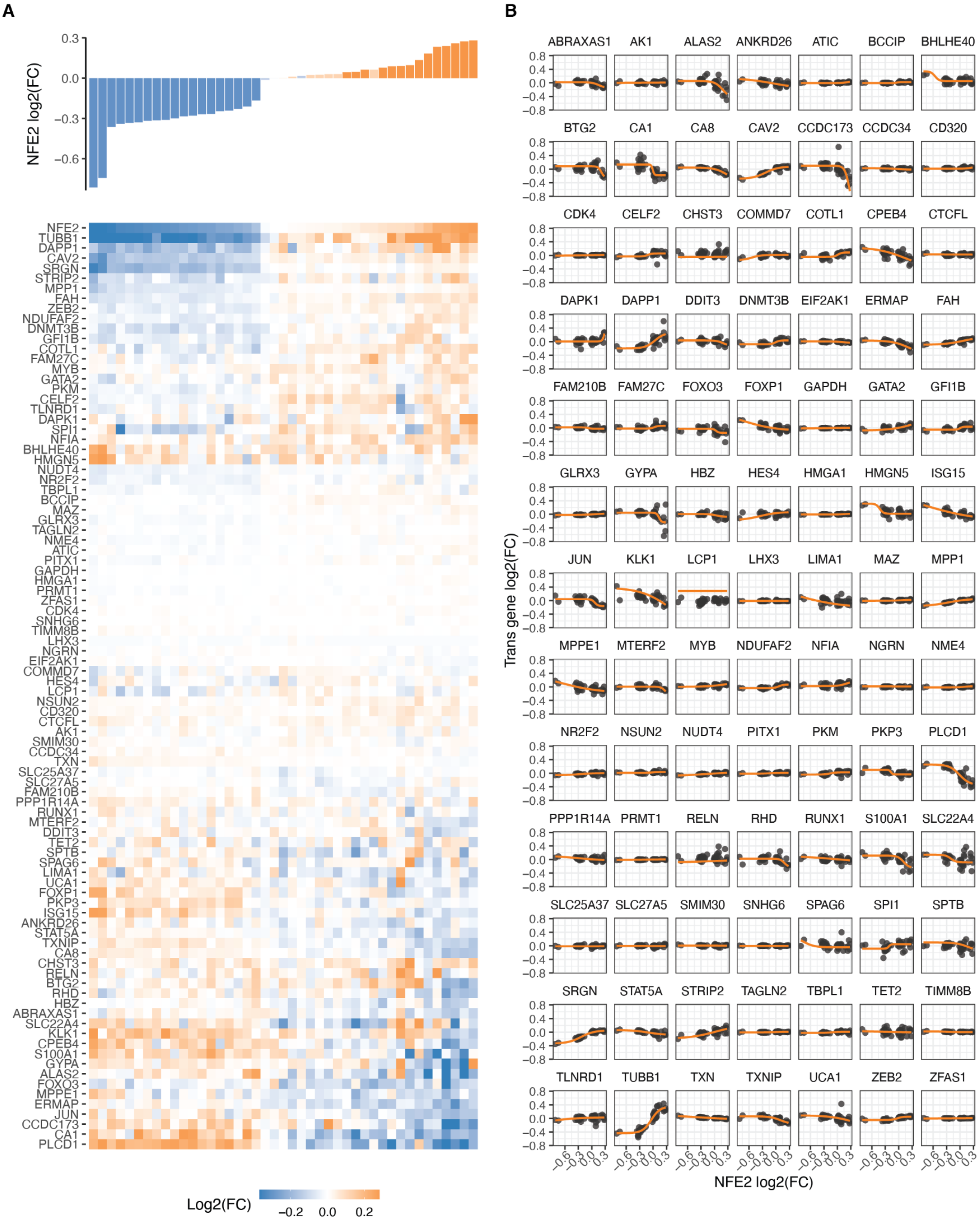


**Figure S7: Trans gene responses to NFE2 dosage modulation**

A. Changes in relative expression of all trans genes (bottom heatmap) in response to NFE2 expression (top barplot) upon each distinct targeted NFE2 sgRNA perturbation. The rows of the heatmap (trans genes) are hierarchically clustered based on their expression fold change linked to alterations in NFE2 dosage.
B. Dosage response curves are plotted for each trans gene against changes in NFE2 expression. The orange line represents the sigmoid model fit.

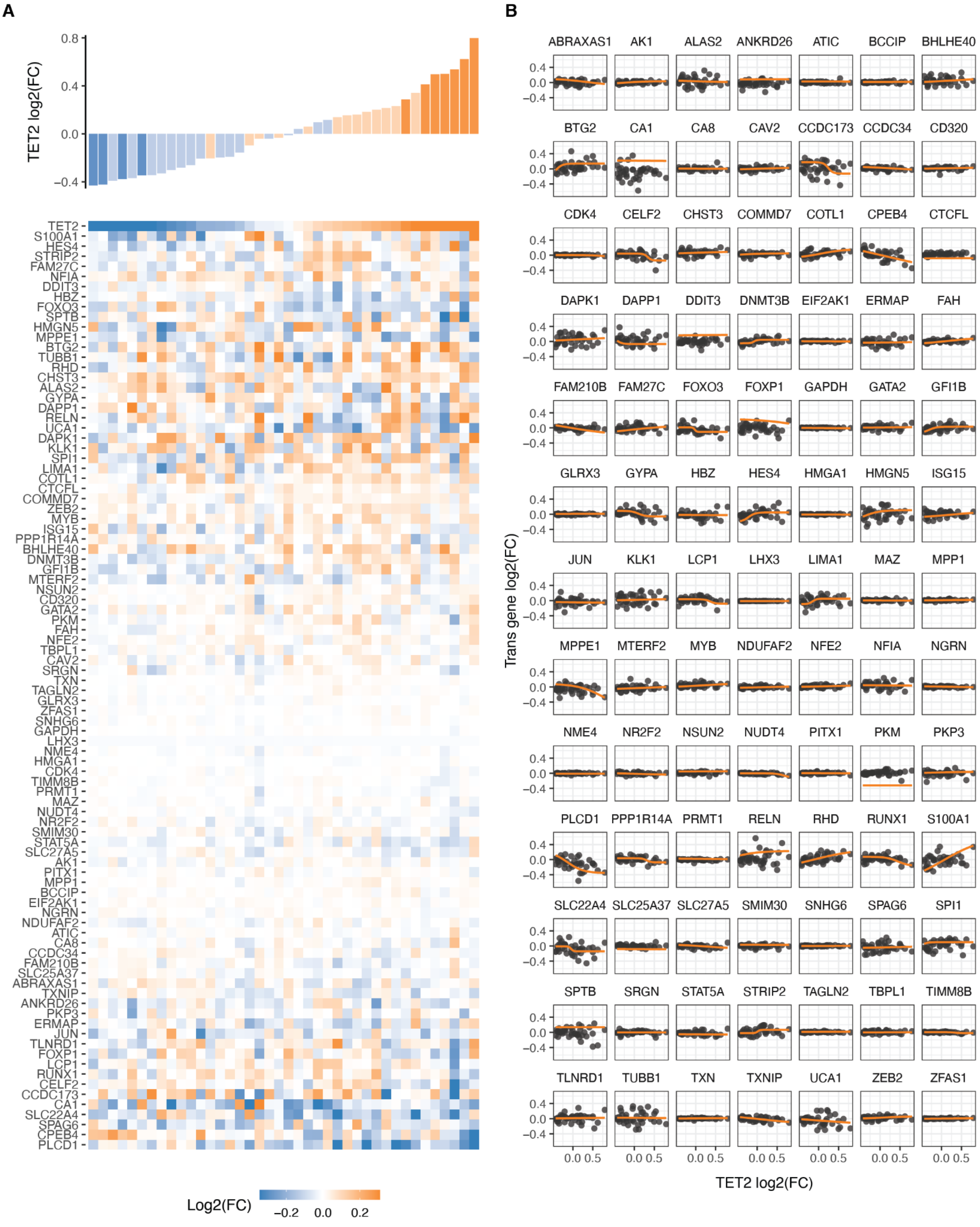


**Figure S8: Trans gene responses to TET2 dosage modulation**

A. Changes in relative expression of all trans genes (bottom heatmap) in response to TET2 expression (top barplot) upon each distinct targeted TET2 sgRNA perturbation. The rows of the heatmap (trans genes) are hierarchically clustered based on their expression fold change linked to alterations in TET2 dosage.
B. Dosage response curves are plotted for each trans gene against changes in TET2 expression. The orange line represents the sigmoid model fit.

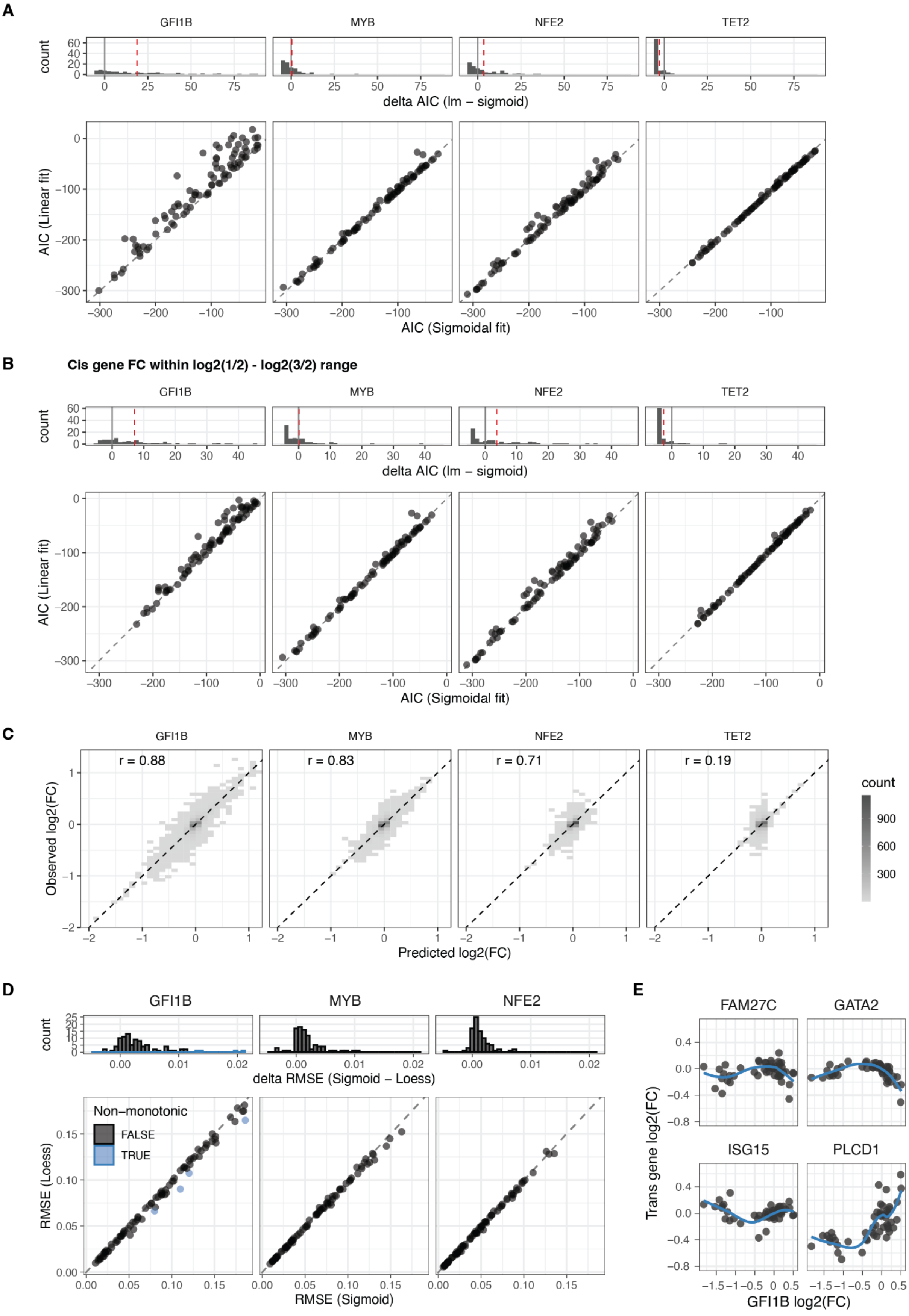


**Figure S9: Dosage response linear and non-linear model fitting**

A. Distribution of the difference in Akaike Information Criterion ($\Delta AIC_{linear\text{-}sigmoid}$) after fitting the sigmoidal or linear model for each trans gene based on the gradual expression perturbations of the four cis genes (top panel), and the direct comparison of the AIC of each fit (bottom panel). Red lines indicate median $\Delta$AIC.
B. Same as A but only fitting the models on those sgRNA perturbations that lead to a cis gene dosage change bounded between log2(1/2) and log2(3/2).
C. Agreement between observed and predicted trans genes expression fold change upon cis gene dosage modulation across a 10-fold cross-validation scheme.
D. Comparison of the Root Mean Square Error (RMSE) of the sigmoid model on the different trans genes dosage responses to the RMSE of the equivalent loess fit (bottom panel). In blue are highlighted the non-monotonic responses that correspond to the top four $\Delta RMSE_{sigmoid\text{-}loess}$ ($RMSE_{sigmoid}$ - $RMSE_{loess}$) values (top panel).

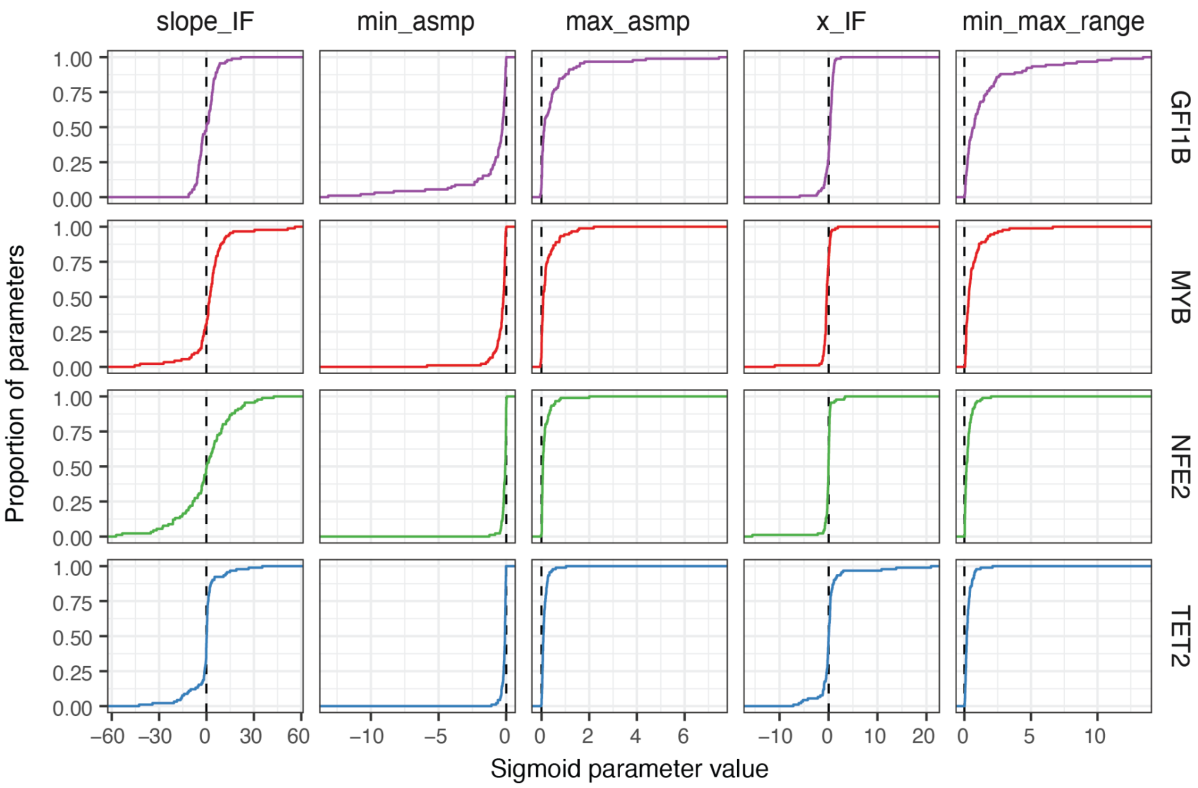


**Figure S10: Distribution of the fitted parameters of sigmoidal model on dosage responses**

Cumulative distribution of the four fitted parameters (first four columns) of the sigmoid model across genes given the independent perturbation of the four TFs (rows). slope_IF = slope of dosage response curve at the inflection point, min_asmp = minimum asymptote (minimum trans gene dosage level), max_asmp = maximum asymptote (maximum trans gene dosage level), x_IF = TF expression FC at the dosage response inflection point.

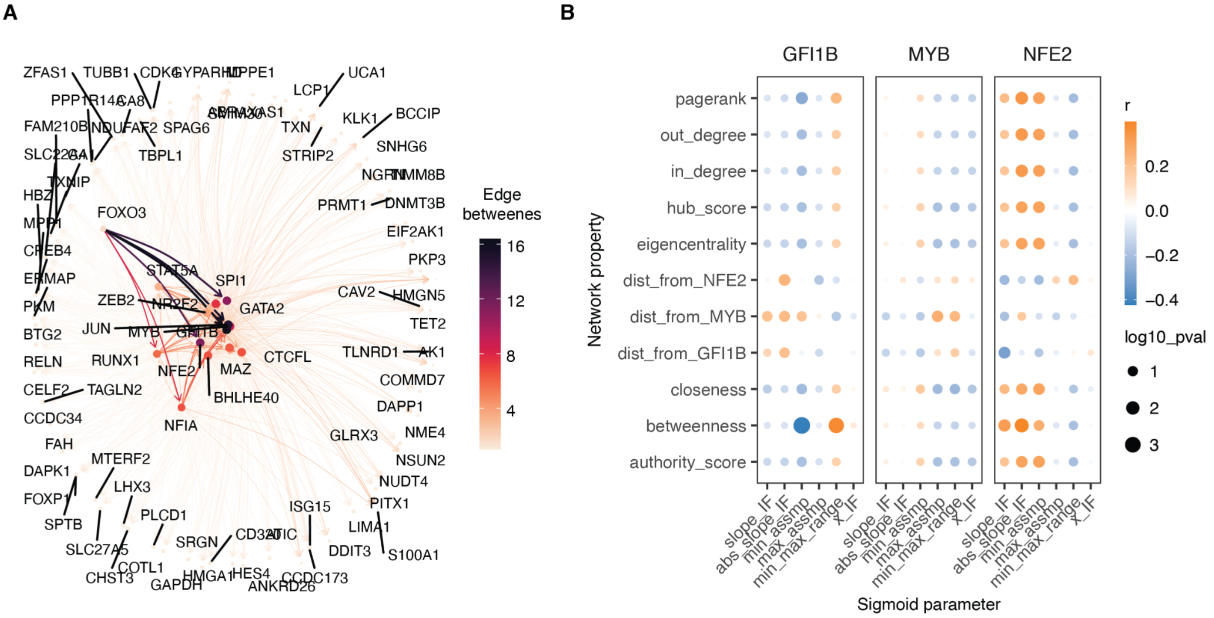


**Figure S11: Relationship gene properties and TF-target network properties with TF dosage responses**

A. A regulatory network constructed based on TF-target gene data Minaeva et al. 2024 with nodes and edges coloured by betweenness. Nodes are sized by their degree.
B. Heatmap illustrating the correlation between the sigmoid parameters in response to cis-gene modulation and network centrality metrics calculated based on the regulatory networks from Minaeva et al. 2024. Point size is scaled to -log10 p-value.

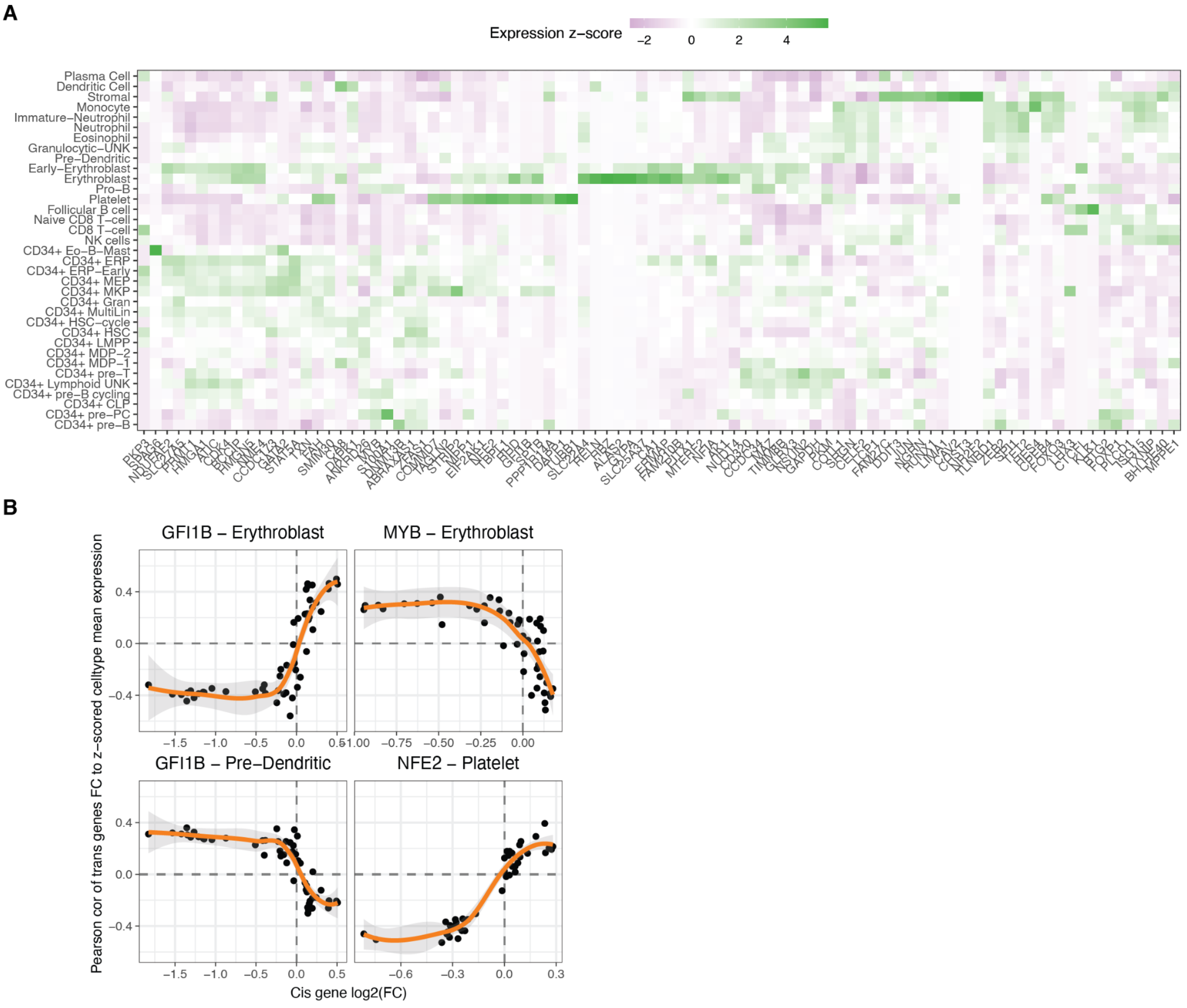


**Figure S12: Transcriptional similarity among bone marrow cell types at different TF dosage levels**

A. Normalised z-score mean expression across donors for targeted genes within each bone marrow cell type (Data from the Human Cell Atlas).
B. Examples of trends of correlation of trans genes expression with the TF change in dosage. The title specifies the cis gene and the cell type for which the trans effects of TF dosage modulation have been contrasted to.
